# An effector protein that protects a fungal pathogen from the plant microbiota during host colonisation

**DOI:** 10.64898/2026.06.10.731356

**Authors:** Liz Florez, Victor Flores-Nuñez, Carolina Francisco, Henry Berndt, Matthias Leippe, Liam Cassidy, Andreas Tholey, Susanne Braun, Graziella Reinhardt, Andrea Sanchez-Vallet, Eva Stukenbrock

## Abstract

*Zymoseptoria tritici* is the causal agent of Septoria tritici blotch, one of the most economically important wheat diseases worldwide. One of the few cloned wheat resistance genes against *Z. tritici*, *Stb6*, recognizes the secreted fungal effector AvrStb6. Although AvrStb6 has been extensively studied as an avirulence determinant, its biological function during host colonization remains unknown. Based on the amphipathic nature of the predicted structure of AvrStb6, we hypothesized the effector to function as a membrane-active antimicrobial protein. However, in vitro growth inhibition assays demonstrated that AvrStb6 does not directly inhibit the growth of wheat-associated bacteria across multiple bacterial genera and experimental conditions. Instead, microbiome analyses of wheat apoplastic fluid revealed shifts in bacterial abundance associated with the presence or absence of *AvrStb6* in a susceptible cultivar (without Stb6-induced resistance). This prompted us to further explore other putative microbiome-related functions of AvrStb6. In vitro confrontation assays further showed that deletion of *AvrStb6* increased the sensitivity of *Z. tritici* to antagonistic wheat-associated bacteria, particularly *Pseudomonas* and *Pantoea* spp. This phenotype was conserved across independent fungal genetic backgrounds and across virulent and avirulent AvrStb6 variants. Fluorescence-based co-culture assays additionally showed reduced fungal growth and increased bacterial proliferation in the absence of *AvrStb6* during interactions with *Pseudomonas* spp., but not with the control bacterium *Escherichia coli*. Finally, biochemical assays demonstrated that AvrStb6 associates with the *Z. tritici* cell wall in vitro, whereas other secreted fungal effectors do not. Collectively, our findings identify a previously uncharacterized role of AvrStb6 in protecting *Z. tritici* from antagonistic wheat-associated bacteria by associating with the fungal cell wall. More broadly, this work highlights that fungal effectors may contribute to microbial competition and ecological adaptation beyond their established roles in host immune recognition.

**Author summary:** Plant pathogens secrete proteins that help them colonize their hosts. Some of these proteins are recognized by plant immune receptors and trigger disease resistance, but their original biological functions often remain unclear. We investigated the role of AvrStb6, a protein produced by the wheat pathogen that causes Septoria tritici blotch. AvrStb6 is best known because it is recognized by a wheat resistance gene, yet its contribution to fungal growth and survival has remained unknown.

We initially tested whether AvrStb6 directly inhibits bacteria that live on wheat leaves, but found no evidence that it acts as an antimicrobial protein. Instead, we discovered that AvrStb6 influences interactions between the pathogen and wheat-associated bacteria. Fungal strains lacking AvrStb6 were more sensitive to several bacterial species that naturally occur in wheat, particularly members of the genera *Pseudomonas* and *Pantoea*. We also found that AvrStb6 can associate with the fungal cell wall, suggesting that it helps protect the pathogen during encounters with antagonistic bacteria.

Our findings reveal an unexpected role for a fungal effector in microbial competition and show that pathogen proteins traditionally studied in the context of plant immunity can also influence interactions with other microbes. This work highlights the importance of considering the broader microbial community when studying plant-pathogen systems.

## Introduction

Plants are continuously exposed to abiotic and biotic stressors and have evolved multilayered defence strategies to limit pathogen invasion. Plant cell surface receptors recognise pathogen-derived molecules and activate signalling pathways associated with defence and adaptation responses (De Falco & Zipfel, 2021; Ngou et al., 2022; Contreras et al., 2023). In parallel, plants recruit beneficial microbes that contribute to disease suppression and pathogen resistance (Ali et al., 2023; Hussain et al., 2023; Du et al., 2025). These plant-associated microbes can inhibit pathogens directly, compete for nutrients and ecological niches, or produce antimicrobial metabolites (Danhorn & Fuqua, 2007; Liu et al., 2017; Rayanoothala et al., 2020; Gu et al., 2020). For example, *Pseudomonas* and *Bacillus* species contribute to disease resistance in the *Arabidopsis* rhizosphere (Kudjordjie et al., 2023), while a *Bacillus* species reduces *Fusarium oxysporum* colonization in cucumber plants (Liu et al., 2017). Together, these observations support the view that the plant microbiome constitutes an external yet integral component of plant immunity that pathogens must overcome during colonization.

Plant pathogens have evolved multiple strategies to suppress host defences and establish infection, including the secretion of effector proteins that manipulate host physiology or immunity. For example, the *Ustilago maydis* effector Cmu1 suppresses salicylic acid biosynthesis in maize, thereby promoting fungal virulence (Djamei et al., 2011), while the *Magnaporthe orzyae* effector SLP1 binds chitin oligosaccharides and competes with the rice chitin receptor, thereby suppressing chitin-triggered immunity Mentlak et al., 2012). Beyond host immune manipulation, fungal effectors are increasingly implicated in interactions with plant-associated microbial communities. In the tomato pathogen *Verticillium dahliae*, several secreted effectors, including VdAve1, VdAMP2, and VdAMP3, display selective antimicrobial activity against plant-associated bacteria and contribute to microbiome manipulation during host colonization (Snelders et al., 2020, 2021, 2023). However, these studies have focused almost exclusively on direct antimicrobial activity as the main mechanism by which effectors influence microbial communities. Additional roles of fungal effectors in mediating interactions with plant-associated microbiota have been hypothesized but remain largely unexplored experimentally (Flores-Nuñez & Stukenbrock, 2024). In this study, we aim to investigate microbe-interacting roles of effector genes in the *Zymoseptoria tritici* - wheat system.

*Z. tritici*, the causal agent of Septoria tritici blotch (STB), colonises the wheat apoplast during an extended asymptomatic phase following stomatal entry (Sánchez-Vallet et al., 2015; Brunner et al., 2019). Throughout this stage of infection, the fungus secretes effector proteins into the apoplastic environment, where they may interact not only with host components but also with wheat-associated microbial communities. To date, molecular studies in the *Z. tritici*–wheat pathosystem have focused primarily on effector-mediated manipulation of host immunity rather than interactions with the plant-associated microbiota. For example, the effector ZtSSP2 interacts with the wheat E3 ubiquitin ligase TaE3UBQ to suppress host-induced cell death (Karki et al., 2021), while LysM effectors bind to chitin in the fungal cell wall to prevent detection and activation of chitin-triggered plant immunity (Tian et al, 2021; Sanchez-Vallet et al, 2020). Only one *Z. tritici* effector (Zt6) has been reported to have cytotoxic activity to plant cells and to exhibit antimicrobial activity against bacteria and yeast (Kettles et al, 2018). In contrast, several other *Z. tritici* effectors have been characterised mainly as avirulence determinants recognized by specific wheat resistance genes, including AvrStb6, Avr3D1, and AvrStb9 (Zhong et al., 2017; Meile et al., 2018; Kema et al., 2018; Amezrou et al., 2023). Although these effectors are central to host recognition, their biological functions during fungal colonization remain poorly understood.

Among these, AvrStb6 is one of the best characterised avirulence effectors in *Z. tritici*. The secreted protein is recognized by the wheat immune receptor Stb6 and transcription of the AvrStb6 gene peaks during early stages of infection, particularly during asymptomatic apoplastic colonization (Siah et al., 2010; Alassimone et al., 2024). Despite widespread deployment of *Stb6* in commercial wheat cultivars, *AvrStb6* remains highly prevalent in field populations, with more than 70 haplotypes identified worldwide (Brunner & McDonald, 2018; Stephens et al., 2021). Although these haplotypes display substantial sequence diversification, premature stop codons and complete gene loss are rare, suggesting strong selective pressure to maintain AvrStb6 function (Sampaio et al, 2025). This observation, together with high gene expression during infection, indicates that AvrStb6 may contribute to fungal fitness beyond its role as an avirulence determinant.

Recent localization studies demonstrated that AvrStb6 is secreted into the wheat apoplast and potentially accumulates around the developing fungal hyphae (Alassimone et al., 2024), suggesting that the protein may interact with components of the apoplastic environment, including the wheat membrane receptor Stb6. Combined with structural predictions and machine-learning analyses, these observations led us to hypothesize that AvrStb6 contributes to interactions between *Z. tritici* and the wheat-associated microbiota. Here, we investigated whether AvrStb6 influences fungal interactions with wheat-associated bacteria and how these interactions affect pathogen colonization. Using microbiome profiling together with in vitro confrontation, fluorescence and biochemical approaches, we show that AvrStb6 reduces the sensitivity of *Z. tritici* to antagonistic apoplastic bacteria through association with the fungal cell wall. These findings provide evidence to a previously uncharacterized function of AvrStb6 and highlight how fungal effectors can contribute to microbial competition within the host environment.

## Materials and methods

### Fungal material

The following *Z. tritici* strains were used in this study: IPO323, IPO323 *ΔAvrStb6*, 3D7-mCherry, 3D7-mCherry *ΔAvrStb6*, and 3D7-mCherry + *AvrStb6*_1E4_ (Kema & van Silfhout, 1997; Meile et al, 2020; Alassimone et al, 2024). Details of all fungal isolates are provided in S1 Table. Cryostocks of the fungal isolates were revived on YMS medium (0.4% [w/v] malt extract, 0.4% [w/v] sucrose, 0.4% [w/v] yeast extract) and incubated at 18°C for five days. Cells were harvested by centrifugation (3,273 *g*, 10 min, room temperature) and resuspended in sterile water to a final concentration of 1 × 10⁶ cells/mL for wheat infection assays or 2 × 10⁷ cells/mL for in vitro confrontation assays. IPO323-derived isolates were used for wheat infection experiments, whereas 3D7-derived strains were used for in vitro fluorescence assays. All *Z. tritici* isolates were included in the in vitro confrontation assays.

### Fungal cell wall stress assays

To assess sensitivity to cell wall and membrane stress, *Z. tritici* strains were grown on YMS agar supplemented with the following stress-inducing compounds (1M Sorbitol, 0.5M NaCl, 500 µg/mL Congo Red, 200 µg/mL CalcoFluor). Fungal cultures were grown in liquid YMS medium for five days at 18°C, harvested by centrifugation (3,273 g, 10 min, room temperature), and resuspended in sterile water to a concentration of 1 × 10⁷ cells/mL. Ten-fold serial dilutions were prepared, and 3 µL of each dilution was spotted onto YMS agar plates containing the indicated compounds. Plates were incubated at 18°C for 4 days and photographed. Growth and colony formation were compared qualitatively between strains to evaluate sensitivity to cell wall and membrane stress. The assay was performed independently three times with similar results

### Wheat infection assays

Fourteen-day old *Triticum aestivum* Obelisk plants were spray-inoculated with water (mock), *Z. tritici* IPO323 wild-type (IPO323), and *Z. tritici AvrStb6* gene deletion mutant (IPO323 *ΔAvrStb6*) isolates. Fungal spray inoculum was adjusted to 1 x 10^6^ cells/mL in sterile water supplemented with 0.1% (v/v) Tween 20. Fungal inoculum was sprayed between 5 to 10 cm distant to the node of the axillary bud of the adaxial side of the third leaf, considering the cotyledon as a first leaf. Post inoculation, plants were maintained in a greenhouse environment and kept with a 16-h light period, 20-22°C temperature for up to three weeks post inoculation. Humidity was kept at 100% for the first two days post inoculation (dpi) and brought down to 80% for the remainder of the experiment.

Disease development was monitored every three days from 3 to 21 dpi. The total number of inoculated leaves per treatment was recorded at the beginning of the experiment and remained constant throughout the assay. At each time point, the number of leaves displaying necrosis or pycnidia was scored independently. Necrosis incidence (%) and pycnidia incidence (%) were calculated as the proportion of symptomatic leaves relative to the total number of inoculated leaves × 100. The experiment was performed independently three times, generating three biological replicates per treatment and time point.

### Wheat apoplast fluid extraction

Apoplastic fluid (AF) was extracted following the protocol described by Francisco et al. (2024, *bioRxiv*). For each treatment, five leaves from independent plants were collected at 4 and 8 dpi. Leaves were washed twice with phosphate-buffered saline (PBS) for 5 min per wash, followed by a wash with PBS containing 0.01% (v/v) Tween 20, and finally rinsed three times with sterile deionized water. Leaves were then submerged in 400 mL of infiltration buffer (5 mM sodium acetate, 0.2 M calcium chloride, pH 4.3) supplemented with two dissolved Complete™ EDTA-free protease inhibitor cocktail tablets (Roche Applied Science, Penzberg, Germany). Vacuum infiltration was performed at 900 mbar for 10 min, after which the vacuum was slowly released over approximately 30 min. This vacuum cycle was repeated two to three times to ensure maximal infiltration of the apoplastic space. The excess buffer was removed by briefly blotting the leaves between sheets of soft paper towel. The leaves were then placed in a 20 mL blunt-end needleless syringe (without the plunger), which was inserted into a 50 mL Falcon tube and centrifuged at 2000 *g* at 4°C for 4 min. The centrifugation step was repeated up to three times to maximize recovery of apoplastic fluid. Two 500 µL aliquots of AF were lyophilized overnight and subsequently resuspended in 200 µL of 100 mM ammonium acetate (pH 7.0) before storage at −20°C for Western blotting or proteomic analysis. In addition, 1 mL of AF was plated on different media for microbial isolation.

### Isolation and establishment of a microbial culture collection from wheat apoplastic fluid

From one mL of freshly extracted AF, a culture collection representing microorganisms enriched in the intercellular space of wheat leaves was established. AF samples from all three experimental conditions (mock, IPO323, and IPO323 *ΔAvrStb6*) were used for culture collection establishment. To maximize microbial recovery, 100 µL of either undiluted or 1:10 diluted AF was plated on three media: Trypticase Soy agar (TSA; 7.5 g/L BBL Trypticase Soy Broth, Becton Dickinson, NJ, USA; 15 g/L agar) and ‘*Pseudomonas syringae*’ medium (PSM; 15 g/L sucrose, 3.75 g/L peptone, 0.38 g/L K₂HPO₄, 0.1 g/L MgSO₄, 15 g/L agar) as non-selective bacterial media; and Reasoner’s 2A agar (R2A; 18.1 g/L R2A agar, Carl Roth, Karlsruhe, Germany) for the isolation of slow-growing bacteria. Three technical replicate plates were prepared for each treatment–medium combination. Plates were incubated at 22°C for four days or until visible colonies appeared. For microbial isolation, colonies displaying distinct morphotypes (e.g., differences in size, shape, texture, or pigmentation) were transferred to fresh agar plates and incubated again for four days at 22°C. This subculturing step was repeated twice to obtain pure cultures. Individualized colonies were then transferred to culture tubes containing 25 mL of Trypticase Soy Broth (TSB) and incubated at 22°C, 145 rpm for seven days. Cryostocks were prepared by mixing 900 µL of liquid culture with 1 mL of 70% (v/v) YMS glycerol medium and stored for long-term preservation.

### 16S rRNA-based identification of wheat apoplastic bacteria

For DNA extraction, 100 µL of liquid culture from bacterial isolates recovered from wheat apoplastic fluid was mixed with 50 µL TE lysis buffer (10 mM Tris-HCl, 1 mM EDTA, pH 8.0, supplemented with 0.1% Triton X-100) and incubated at 99°C for 10 min to lyse cells. Samples were subsequently centrifuged at 13,000 *g* for 10 min, and the supernatant containing genomic DNA was transferred to a new tube and used as template for PCR amplification.

To identify bacterial isolates, partial regions of the 16S rRNA gene were amplified as previously described (Marchesi et al., 1998; Bellemain et al., 2010). The bacterial 16S rRNA V3–V7 region was amplified using primers 341F (5′-CCTACGGGAGGCAGC-3′) and 1192R (5′-ACGTCATCCCCACCT-3′) under the following cycling conditions: initial denaturation at 94°C for 2 min; 25 cycles of 94°C for 30 s, 57°C for 30 s, and 72°C for 1 min; followed by a final extension at 72°C for 10 min. PCR product size was verified by agarose gel electrophoresis. Confirmed PCR products were submitted for Sanger sequencing at the Institute of Clinical Molecular Biology (Kiel, Germany) using both forward and reverse primers.

Resulting AB1 files were processed using the IsolateR package (v1.0) to generate trimmed high-quality sequences and perform taxonomic assignment using default parameters. Because only partial 16S rRNA sequences were obtained, taxonomic classification was considered reliable primarily at the genus level. Differential abundance analysis was performed in R using relative abundance values at the genus level. Log2 fold change values were calculated as follows: log2FC = log2((relative abundance in IPO323 + 1e-6)/(relative abundance in IPO323 Δ*AvrStb6* + 1e-6)), where a pseudocount (1e-6) was added to avoid division by zero when a genus was absent from one treatment. Positive log2FC values indicated higher relative abundance in IPO323, whereas negative values indicated higher relative abundance in IPO323 Δ*AvrStb6*.

### Metabarcoding of whole leaf-associated microbial communities

For microbiome profiling, leaves from five wheat plants per treatment were collected at 4 and 8 dpi and processed for 16S rRNA V4 amplicon sequencing. Each plant was considered an independent biological replicate. Sample processing and DNA extraction were performed as described by Seybold et al. (2021) with modifications. Briefly, loosely attached particles were removed from the leaf surface, after which leaves were dried and ground in liquid nitrogen using a mortar and pestle. The metagenomic DNA of grinded leaf tissue was extracted using FastDNA™ Spin Kit for Soil (MP Biomedicals) with the modifications reported by (Seybold et al., 2020). The DNA samples were sent to the University of Minnesota Genomics Center (UMGC), where the metabarcoding sequencing of the 16S-rRNA-V4 (515F/816R primers) was performed as described in Flores-Nunez, et al. 2026. Peptide nucleic acid (PNA) clamps were added in the reactions to reduce the amplification of plant chloroplast and mitochondrial DNA (Lundberg et al, 2013). Paired-end 2 × 300 bp sequencing was performed with an Illumina MiSeq v3 instrument. Data was demultiplexed, resulting in 30 libraries The generated raw reads have been deposited under the SRA accession number PRJNA1474436. Raw sequencing data were processed as described by Flores-Núñez et al. (2026). After quality filtering and Operational Taxonomic Unit (OTU) processing, the dataset comprised 25 samples from three treatments collected at 4 and 8 dpi, with a median sequencing depth of 779 reads per sample and a total of 1313 OTUs. All downstream microbiome analyses and data visualization were performed in R (v4.1.1; R Core Team, 2021).

Alpha diversity analyses were conducted using the vegan package, while plots were generated using ggplot2 (v4.0.1) and ggpubr (v0.6.3; Wickham, 2012). To account for variation in sequencing depth, the OTU table was rarefied to the minimum read count (394 reads per sample). Alpha diversity within samples was estimated using the Shannon diversity index, and differences between infection treatments were assessed using a Kruskal–Wallis rank-sum test followed by Dunn’s post hoc test.

Beta diversity was estimated using Bray–Curtis dissimilarity and visualized using non-metric multidimensional scaling (NMDS). The effect of every treatment on community composition was evaluated by PERMANOVA using the model Bray–Curtis ∼ treatment. Relative abundance values were calculated and visualized at the genus level. Differential abundance analysis was performed using the metagenomeSeq package (v1.52.0; Paulson et al., 2013) using cumulative sum scaling normalization. Only OTUs with ≥2 reads per sample were retained for analysis, and OTUs with a p-value <0.05 and prevalence >50% across samples were considered differentially abundant.

### *Pichia pastoris* protein expression and purification

The *Pichia pastoris* protein expression system was used to produce AvrStb6 protein variants fused to an N-terminal 6×His tag followed by an enterokinase (EK) cleavage site. Synthetic DNA fragments encoding His-EK-AvrStb6_1E4_ and His-EK-AvrStb6_1A5_ were cloned into the pPICZ**α** vector under the control of the inducible AOX1 promoter and fused to the **α**-factor secretion signal to enable extracellular secretion of the recombinant proteins. The constructs were transformed into the wild-type *Pichia pastoris* X-33 strain. Transformation was performed by electroporation using 80 µL of *P. pastoris* electrocompetent cells and 10 µg of linearized plasmid DNA. Transformed cells were recovered in 1 mL ice-cold 1 M sorbitol and incubated at 28°C for 1–2 h, and plated onto YPD (Yeast Peptone Dextrose) agar supplemented with 40% (v/v) glucose and 500 µg/mL of Zeocin and incubated at 28°C for 2–3 days until colonies appeared. Candidate transformants were subsequently plated on YPD containing 1000 µg/mL of Zeocin. After two to three days, colonies showing the strongest growth were selected.

For protein production, transformants were streaked onto YPD plates containing 100 µg/mL of Zeocin and incubated for 2 days at 28°C. To generate a preculture, a single colony was inoculated in 3 mL YMS medium and incubated at 28°C with shaking (250 rpm) overnight (16-18 h). Subsequently, 1 mL of preculture was transferred into 250 mL of YMS medium and grown at 28°C, 250 rpm until reaching exponential phase (OD_600_ ≈ 2–6). Cultures were harvested by centrifugation (3000 *g* at room temperature for 5 min), and the pellets were resuspended in 1 L of buffered methanol-complex medium (BMMY) to an OD_600_ of approximately 1. Cultures were incubated at 28°C with shaking (265 rpm). After 24 h of growth, methanol was added to a final concentration of 0.5% to induce gene expression from the AOX1 promoter, and cultures were incubated for an additional 5 h.

Culture supernatants containing secreted recombinant proteins were harvested by centrifugation (13,689 *g*, 4°C, 4 min), and the cleared supernatant was collected and kept on ice. Recombinant proteins were purified by immobilized metal affinity chromatography using His/Ni-NTA resin. The resin was equilibrated with an equilibration buffer and added to the culture supernatant, followed by incubation with gentle mixing to allow binding of His-tagged proteins. After incubation, the resin was collected by centrifugation and transferred to fresh tubes. The beads were washed three times with a wash buffer to remove non-specifically bound proteins, and bound proteins were subsequently eluted using imidazole (250mM) - containing elution buffer (pH 7.4). Elution fractions were collected separately and protein concentrations were determined by measuring the 280nm absorption using a microvolume spectrophotometer (NanoDrop). Protein purification was verified by SDS-PAGE and the product identified by Western blot analysis using an anti-His tg antibody.

To obtain the natural proteoforms of the AvrStb6 variants, the His-tagged proteins were desalted into enterokinase cleavage buffer, digested with enterokinase, and subsequently purified by reversed-phase FPLC using a Hamilton PRP-3 resin and a linear gradient from 0.1% trifluoroacetic acid to 80% acetonitrile in 0.1% trifluoroacetic acid using an ÄKTA Purified FPLC System. Eluted peptide fractions were pooled and lyophilized.

### Western Blot

Protein samples were separated by SDS-PAGE and transferred onto PVDF membranes using a wet transfer system. Prior to transfer, gels and filter papers were equilibrated in cold transfer buffer (25 mM Tris, 40 mM glycine, 2 mM SDS, 20% methanol), and PVDF membranes were activated in methanol and subsequently washed in transfer buffer. Membranes were blocked at 4°C in 5% milk prepared in TBST overnight and incubated with mouse anti-His primary antibody (1:2,000; Agrisera, AS111771) for >1 h at 4°C with shaking, followed by incubation with HRP-conjugated rabbit anti-mouse IgG secondary antibody (1:20,000, Agrisera, AS09627).

### *In vitro* microbial growth assay

For the bacterial growth inhibition assay, bacterial isolates from the *T. aestivum* Obelisk AF culture collection (S2 Table) were used. Cryostocks of the bacterial isolates were revived on Lysogeny Broth (LB) agar plates and incubated at 23°C overnight (16-18 h). Single colonies were selected and grown in LB liquid media at 23°C and shaking (180 rpm) overnight. Overnight cultures were resuspended to an optical density 600 (OD_600_) of 0.02. For the protein-bacteria growth assays, a serial dilution (24µM to 0.047 µM) of AvrStb6_1E4_ and AvrStb6_1A5_ recombinant proteins were resuspended in 10mM sodium phosphate (pH 5.8). Then 90 µL of diluted protein and 10 µL of bacteria isolate (OD_600_ 0.02) were aliquoted per well in 96-well flat-bottom culture plates. Two technical replicates were prepared per bacterial isolate and per protein concentration. *Escherichia coli* strain K12 and the peptide melittin were used as a positive control for bacterial growth inhibition. Plates were incubated in a Synergy H1 plate reader (BioTek, VT, USA) at 23°C with orbital shaking every 15 min (10 s at 300 rpm). For the curve growth assay, bacterial OD_600_ was measured every 15 min after shaking, and for the microsusceptibility assay, bacterial OD_600_ was measured after 24 hours.

### In vitro confrontation assay

Bacterial isolates from the *T. aestivum* Obelisk AF culture collection (S2 Table) were confronted *in vitro* with *Z. tritici* IPO323-and 3D7-related isolates in a one-to-one interaction. Fungal spore suspensions were mixed in melted 0.5x-Fries3 media for a final concentration of 1 x 10^6^ spores/mL. Once solidified, one µL of OD_600_ 0.01 per bacterial isolates was pipetted on top. Per plate, a fungal isolate was confronted with three bacteria isolates from the same genera, having three technical replicates per bacteria isolate. One µL of Hygromycin was added as a positive control for inhibition. Plates were incubated at 23°C for four days. Colony diameter and inhibition halo diameter were measured at four days post incubation. Inhibition halo length was then calculated by subtracting the colony diameter from the inhibition halo diameter to account for differences in bacterial colony size.

### dTomato-fluorescence tagged bacteria design

Selected wheat-associated *Pseudomonas brassicacearum* isolates (S2 Table) were transformed with the dTomato-expressing plasmid pWT-dTomato, kindly provided by Tal Dagan. Electrocompetent cells were prepared from exponentially growing cultures (OD_600_ 0.5) grown in LB medium at 28°C with shaking. Cells were washed three times with ice-cold 300 mM sucrose and resuspended in 300 mM sucrose. Approximately 80 µL of competent cells were mixed with plasmid DNA and transferred into pre-chilled 1 mm electroporation cuvettes. Electroporation was performed using a Gene Pulser system (Bio-Rad) at 1.8 kV, 25 µF, and 200 Ω. Following pulsing, cells were recovered in SOC medium at 28°C with shaking (300 rpm) for 1 hour and plated onto dYT agar supplemented with kanamycin (500 µg/mL). Fluorescent transformants were identified by dTomato fluorescence and integration was verified by sequencing. Transformants were maintained under antibiotic selection. *Escherichia coli* strains expressing dTomato fluorescence were generated using the same procedure.

### In vitro fluorescence assay

For fungal-bacterial fluorescence assays, mCherry-tagged *Z. tritici* isolates (S1 Table) and dTomato-tagged *P. brassicacearum* isolates were used. Seven-day-old *Z. tritici* cultures were resuspended to a spore concentration of 10^6 cells/mL, while overnight-grown *P. brassicacearum* cultures were adjusted to an OD_600_ of 0.02. For each co-culture, 100 µL of fungal suspension and 100 µL of bacterial suspension were aliquoted into individual wells of 96-well plates. Two technical replicates were prepared per fungal-bacterial combination. mCherry-tagged and non-fluorescent 3D7 strains were used as positive and negative controls for mCherry fluorescence, respectively. Similarly, dTomato-tagged and non-fluorescent *E. coli* Top10 strains were used as positive and negative controls for dTomato fluorescence. Plates were incubated in a Synergy H1 plate reader (BioTek, VT, USA) at 23°C with orbital shaking every 15 min (10 s at 300 rpm). dTomato fluorescence was measured using excitation and emission wavelengths of 550 nm and 590 nm, respectively, while mCherry fluorescence was measured using excitation and emission wavelengths of 580 nm and 620 nm, respectively. Fluorescence gain was set to 150 for both channels. mCherry and dTomato fluorescence signals were recorded every 15 min for 48 h.

### Statistical analysis

Data from *in vitro* growth-and fluorescence assays was processed in RStudio (V 2025.09.0). Prior to statistical analyses, a normal distribution of the data was assessed using the Shapiro-Wilk test. Student T test and Analysis of variance (ANOVA) were carried out for two-treatment and multiple-treatment comparisons, respectively, when the assumptions of a normal distribution were met. In ANOVA, if statistical significance was observed (p value < 0.01), a post-hoc Tukey test was used for specific treatment vs treatment comparisons.

### Structural prediction and electrostatic surface analysis

The three-dimensional structure of AvrStb6 was predicted using AlphaFold3 server (Jumper et al, 2021; Abramson et al, 2024). The putative signal peptide (predicted with InterPro v 108.0) was removed before modelling. The most confident model (highest pLDDT score) was further explored in ChimeraX and the PDB files submitted across multiple servers. Structural similarity searches were performed using FoldSeek (van Kempen et al., 2024) and the DALI server (Holm, 2022) to identify potential structural homologs in the Protein Data Bank. Antimicrobial-associated properties were predicted using AMAPEC v1.0 (Mesny & Thomma, 2024; Mesny, 2026) through the Google Colab implementation available at AMAPEC Google Colab Notebook. Electrostatic surface potentials were calculated using the Poisson–Boltzmann method implemented in the PDB2PQR/APBS software suite (Dolinsky et al., 2004; Jurrus et al., 2018) and visualized in ChimeraX.

### Cell-wall binding assay

Alcohol-insoluble residue (AIR) was prepared from *Z. tritici* blastospores grown for 6 d in YPD medium (1% yeast extract, 2% peptone, 2% dextrose) supplemented with 50 µg/mL kanamycin sulfate at 18°C and shaking at 120 rpm. Fungal cells were harvested by centrifugation (3,273 *g*, 4°C, 15 min), washed twice with distilled water, frozen in liquid nitrogen, ground to a fine powder, and lyophilized overnight. The dried material was sequentially extracted with 1:1 (v/v) methanol:chloroform (three times; 30 mL/g, including one overnight incubation), acetone (30 mL/g), 70% ethanol (twice; 30 mL/g, including one overnight incubation), and a final acetone wash (30 mL/g). All extraction steps were performed at 4°C with rotation, followed by centrifugation at 3,273 *g* for 10 min. After the final wash, samples were air-dried in a fume hood overnight, and the resulting dried material was used as AIR for cell-wall binding assays.

For cell-wall binding assays, AIR preparations were incubated separately with individual recombinant His-tagged proteins (S3 Table) at 4°C with shaking (100 rpm) for 1 h, and centrifuged at 13,000 *g* at 4°C for 5 min. Pellets and supernatant fractions were separated, and pellets were resuspended in 100 µL water. Samples were boiled at 95°C for 10 min prior to SDS-PAGE and Western blot analysis as described in the western blot section.

## Results & Discussion

### AvrStb6 in silico predictions

Previous studies have demonstrated that the secreted effector protein AvrStb6 plays a major role in the interaction between *Z. tritici* and wheat cultivars expressing the immune receptor Stb6 (Zhong et al, 2017; Saintenac et al, 2018). Despite its importance in host recognition and virulence, the biological function of AvrStb6 has remained unknown. To gain insight into its potential molecular activity, we firstly performed structural and computational analyses of the mature AvrStb6 protein.

AlphaFold3 (Abramson et al, 2024) predicted AvrStb6 (the allele of the reference isolate IPO323 and the Swiss 1E4 isolate) to form a compact globular structure with relatively high overall confidence (pTM = 0.79) (Fig 1A). The core of the protein exhibited high per-residue confidence scores, whereas reduced confidence was largely confined to flexible surface-exposed and terminal regions. Structural similarity searches using FoldSeek (van Kempen et al., 2024) and DALI (Holm, 2022) did not identify significant structural homologs, whereas the machine learning-based tool AMAPEC v1.0 (Mesny & Thomma, 2024; Mesny, 2026) predicted AvrStb6 to have antimicrobial properties with high confidence (probability = 0.997). Electrostatic surface analysis based on Poisson–Boltzmann calculations revealed spatially separated positively-charged surface patches (regions in blue) together with exposed hydrophobic regions on the predicted AvrStb6 structure (Fig 1B and 1C). This amphipathic surface organization is commonly associated with membrane-and surface-interacting proteins, where electrostatic interactions with phospholipid head groups and hydrophobic contacts can contribute to membrane association (Whited & Johs, 2015: Petti et al, 2026). Together, these analyses suggested that AvrStb6 possesses structural features consistent with potential membrane-or surface-interacting activity and motivated us to perform subsequent functional characterization.

**Figure 1.**
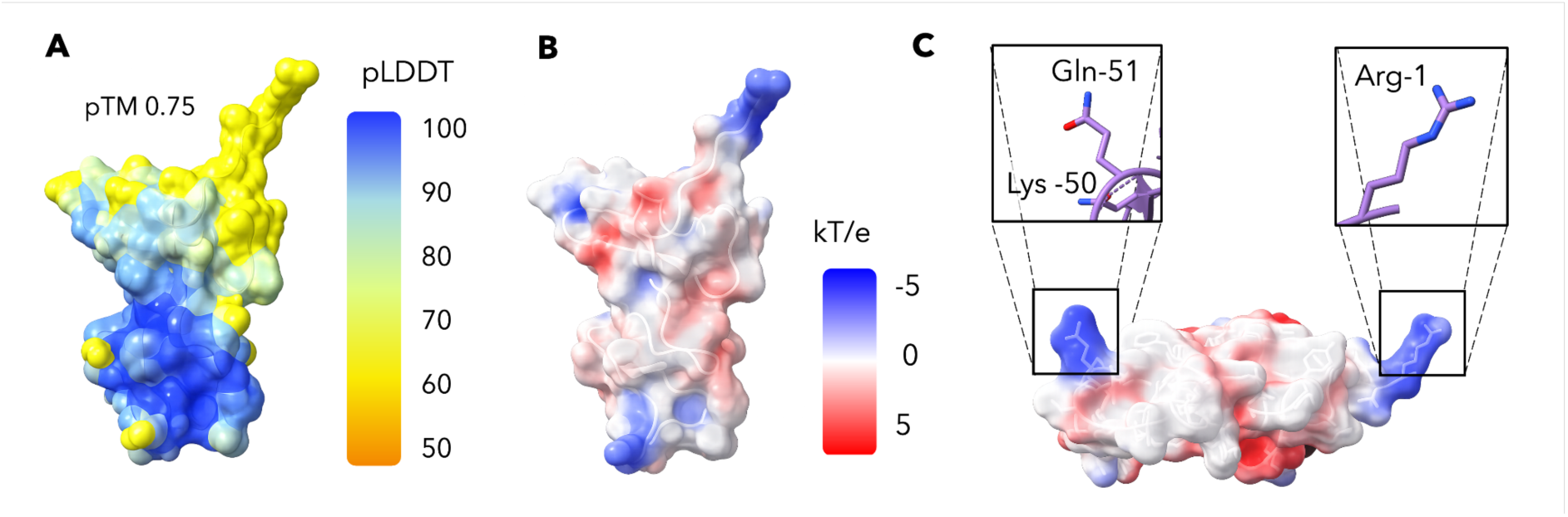
Predicted structure and surface properties of AvrStb6. (A) AlphaFold3-predicted structure of mature AvrStb6 colored according to per-residue confidence (pLDDT). The predicted template modelling (pTM) score is indicated. (B) Poisson–Boltzmann electrostatic surface representation of AvrStb6 showing heterogeneous surface charge distribution (in red) and distinct positively charged surface patches (in blue). (C) Surface-exposed positively charged regions of AvrStb6. Insets highlight representative solvent-accessible residues associated with positively charged surface patches.

### AvrStb6 contributes modestly to virulence but strongly alters the wheat microbiome

To investigate whether the predicted structural properties of AvrStb6 contribute to fungal pathogenicity, we first assessed the role of the effector during wheat infection using the cultivar Obelisk, which lacks the cognate resistance gene Stb6. Wheat plants were inoculated with the *Z. tritici* isolate IPO323, which expresses AvrStb6 during early host colonization, or with an isogenic *ΔAvrStb6* deletion mutant (Kema et al., 2018). Deletion of *AvrStb6* resulted in a modest reduction in disease development, with decreased pycnidia formation (20-28% reduction, ANOVA, p < 0.01) and diminished host necrosis (20-25% reduction, ANOVA, p < 0.01) relative to the IPO323 wild type (Fig 2A, 2B and 2D). In contrast, both strains displayed comparable growth in vitro (S1 Fig), indicating that the reduced virulence phenotype was not associated with impaired fungal growth under laboratory conditions, but related to infection-associated traits of the fungus.

**Figure 2.**
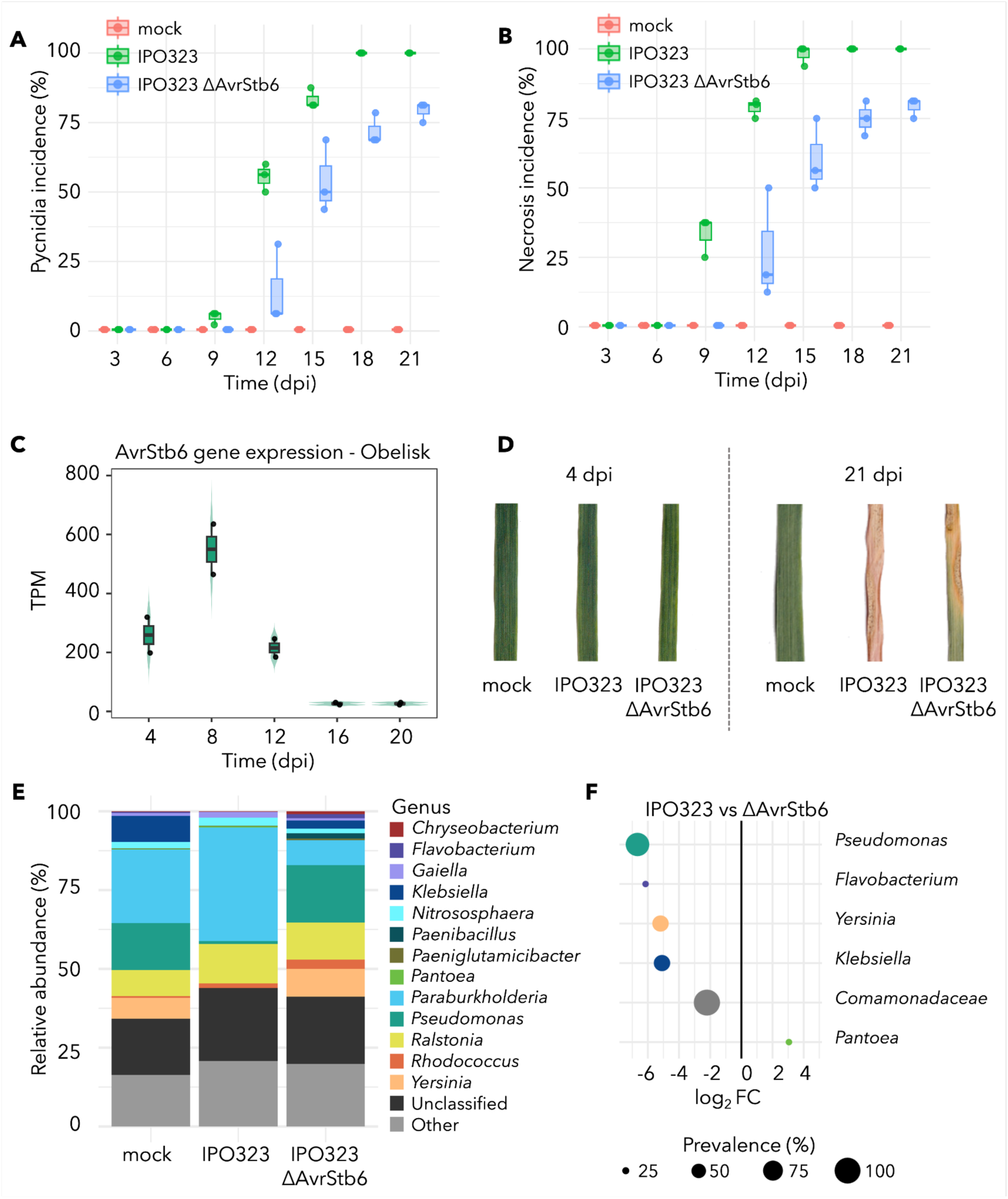
AvrStb6 modestly affects disease symptoms but drives major changes in wheat leaf microbiome composition. Wheat cultivar Obelisk (lacking Stb6) was inoculated with water (mock), *Zymoseptoria tritici* IPO323 (wild type), or the IPO323 *ΔAvrStb6* deletion mutant. (A) Pycnidia incidence (%) and (B) necrosis incidence (%) over days post-inoculation (dpi). (C) *AvrStb6* expression (TPM) in infected leaves across the disease time course. (D) Representative leaf phenotypes at early (4 dpi) and late (21 dpi) infection stages. (E) Relative abundance (%) of bacterial genera based on 16S rRNA gene profiling of whole-leaf communities at 4 dpi. (F) Differential abundance analysis comparing IPO323 and IPO323 *ΔAvrStb6* bacteria composition. A differential abundance analysis was performed using the metagenomeSeq package (v 1,52,0; Paulson et al, 2013) using the cumulative sum scaled OTU table. Log₂ fold change indicates enrichment or depletion in IPO323 relative to IPO323 *ΔAvrStb6*; point size reflects prevalence across samples.

Given the predicted membrane-interacting and antimicrobial-properties of AvrStb6, we hypothesized that the effector may contribute to virulence by altering the composition of wheat-associated bacterial communities during infection. To investigate this possibility, we first determined infection stages associated with elevated AvrStb6 gene expression in planta in order to guide microbiome sampling. Consistent with previous gene expression studies showing that *AvrStb6* expression peaks during the transition from biotrophic to necrotrophic growth (Rudd et al., 2015; Zhong et al., 2017; Kema et al., 2018), expression analysis in Obelisk revealed increased AvrStb6 transcript accumulation from mid-biotrophy to early necrotrophy (Fig 2C). Based on this expression pattern, 16S rRNA amplicon profiling was performed at 4 and 8 dpi.

Bacterial community composition of wheat leaves was driven by the infection treatment (PERMANOVA, r^2^ = 0.34, p = 0.001 at 4dpi and r^2^ = 0.33, p = 0.016 at 8dpi), where tighter clustering of biological replicates happening at 4 dpi and greater dispersion at 8 dpi (S2 Fig). Alpha diversity measurements were comparable across treatments at 4 dpi but diverged at 8 dpi, where infected samples showed a reduced Shannon diversity tendency relative to mock-treated plants. Given the reduced variability among biological replicates observed at 4 dpi, subsequent analyses focused primarily on this time point. At 4 dpi, bacterial community composition differed between IPO323-and *ΔAvrStb6*-infected leaves (Fig 2E). Most notably, *Pseudomonas* spp. (OTU 23) represented less than 1% of the bacterial community in IPO323-infected plants, compared with approximately 14% and 16% in mock-and *ΔAvrStb6*-treated samples, respectively (Fig 2E). Differential abundance analyses further identified multiple bacterial genera that were significantly affected by the absence of AvrStb6 (Fig 2F). Differential abundance analysis further identified several bacterial taxa associated with the presence or absence of AvrStb6, including *Pseudomonas*, *Flavobacterium*, *Yersinia*, *Klebsiella*, *Comamonadacea* and *Pantoea* (Fig 2F). Among these, *Pseudomonas* spp. displayed the largest and most consistent abundance shift associated with AvrStb6 activity.

To further investigate bacterial populations co-occurring with *Z. tritici* in the wheat apoplast and their putative interaction with AvrStb6, we established a culture collection from apoplastic fluids isolated from Obelisk leaves. In total, 440 bacterial isolates representing 27 genera were recovered. Consistent with the whole-leaf microbiome profiles, *Pseudomonas* spp. were abundant in mock-and ΔAvrStb6-treated samples but were undetectable in the apoplast of IPO323-infected plants (S3C Fig). Differential abundance analyses of the culturable apoplastic fraction additionally identified shifts in *Rahnella*, *Enterobacter*, *Flavobacterium*, and *Dufyella* between treatments (S3D Fig). Together, these results identify *Pseudomonas* spp. as the bacterial group most strongly associated with the presence or absence of AvrStb6 during wheat colonization.

### AvrStb6 does not exert direct antimicrobial activity

Based on the differential abundance of multiple bacterial genera, particularly *Pseudomonas*, in wheat leaves infected with IPO323 (expressing *AvrStb6*) and the IPO323 Δ*AvrStb6* mutant, we hypothesized that AvrStb6 may exhibit antimicrobial activity against wheat-associated bacteria. To test whether AvrStb6 exerts direct bactericidal or bacteriostatic activity, we performed in vitro bacterial growth inhibition assays using the natural proteoform of AvrStb6 protein, obtained by heterologously expression in the yeast *Pichia pastoris*, and bacterial isolates assigned to the genera *Curtobacterium*, *Enterobacter*, *Flavobacterium*, *Pantoea*, *Pseudomonas*, and *Rahnella* following the partial 16S rRNA taxonomic classification (S2 Table). These genera were selected based on their differential abundance in the wheat apoplastic fluid fraction associated with the presence or absence of *AvrStb6*.

At 8 µM, recombinant AvrStb6 protein did not inhibit growth of any tested bacterial strain. In contrast, the positive control, the honeybee membranolytic peptide melittin, completely inhibited bacterial growth at the same concentration (Fig 3A). To assess potential concentration-dependent effects, we next performed minimum inhibitory concentration (MIC) assays using AvrStb6 concentrations up to 24 µM. Whereas melittin displayed a MIC of 1.5 µM against all tested bacteria, AvrStb6 failed to inhibit bacterial growth even at the highest concentration tested (Fig 3B). Together, these results indicate that AvrStb6 alone does not exhibit detectable antimicrobial activity against the selected Gram-negative and Gram-positive wheat-associated bacteria under the tested in vitro conditions.

**Figure 3.**
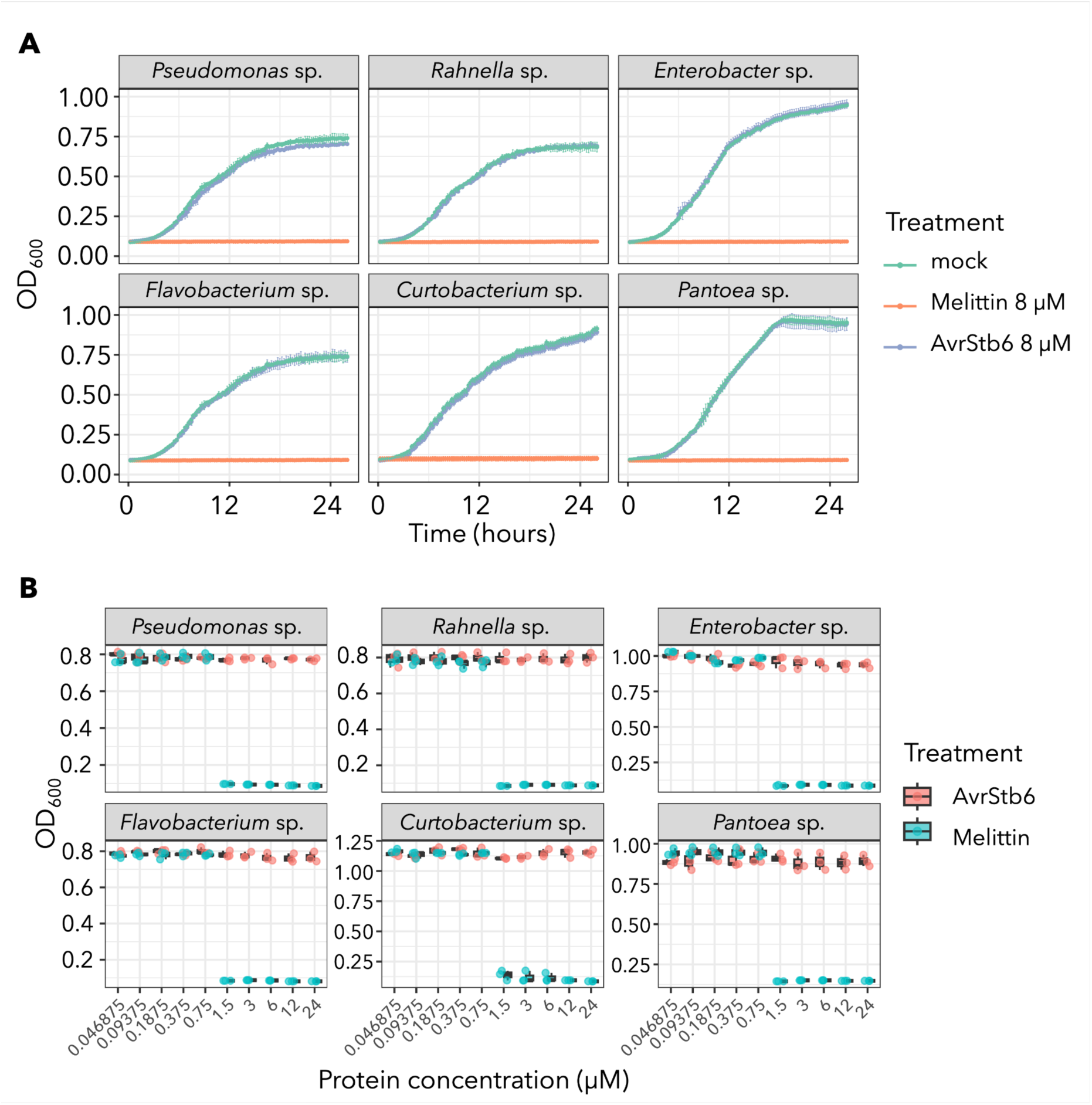
AvrStb6 has no antibacterial activity towards wheat apoplast-associated bacteria. (A) Growth curves (OD₆₀₀) of wheat apoplast-associated bacteria over 24 h. Mock (ultra pure water) served as a non-treatment control and Melittin (honeybee antimicrobial peptide) was used as a positive control for bacterial growth inhibition. (B) Final OD₆₀₀ values were measured at 24 h after exposure of bacteria with different concentrations of AvrStb6 protein (up to 24 µM).

### AvrStb6 reduces sensitivity of *Zymoseptoria tritici* to bacterial antagonism

To further investigate whether the observed differential abundance of bacterial taxa in IPO323-and IPO323 Δ*AvrStb6*-infected leaves could reflect direct bacterial-fungal antagonism, we examined interactions between the IPO323 wild-type and Δ*AvrStb6* mutant *Z. tritici* strains and the same bacterial isolates used in the antimicrobial assays described above. Three isolates from each of the genera *Pseudomonas*, *Rahnella*, *Enterobacter*, *Flavobacterium*, *Curtobacterium*, and *Pantoea* were included in the analysis (S2 Table). In vitro confrontation assays were performed by embedding fungal spores and pseudohyphae in agar followed by application of bacterial inocula onto the agar surface. Most bacterial isolates inhibited *Z. tritici* growth, producing inhibition zones ranging from approximately 1 mm to 25 mm after 4 days of incubation (Fig 4).

**Figure 4.**
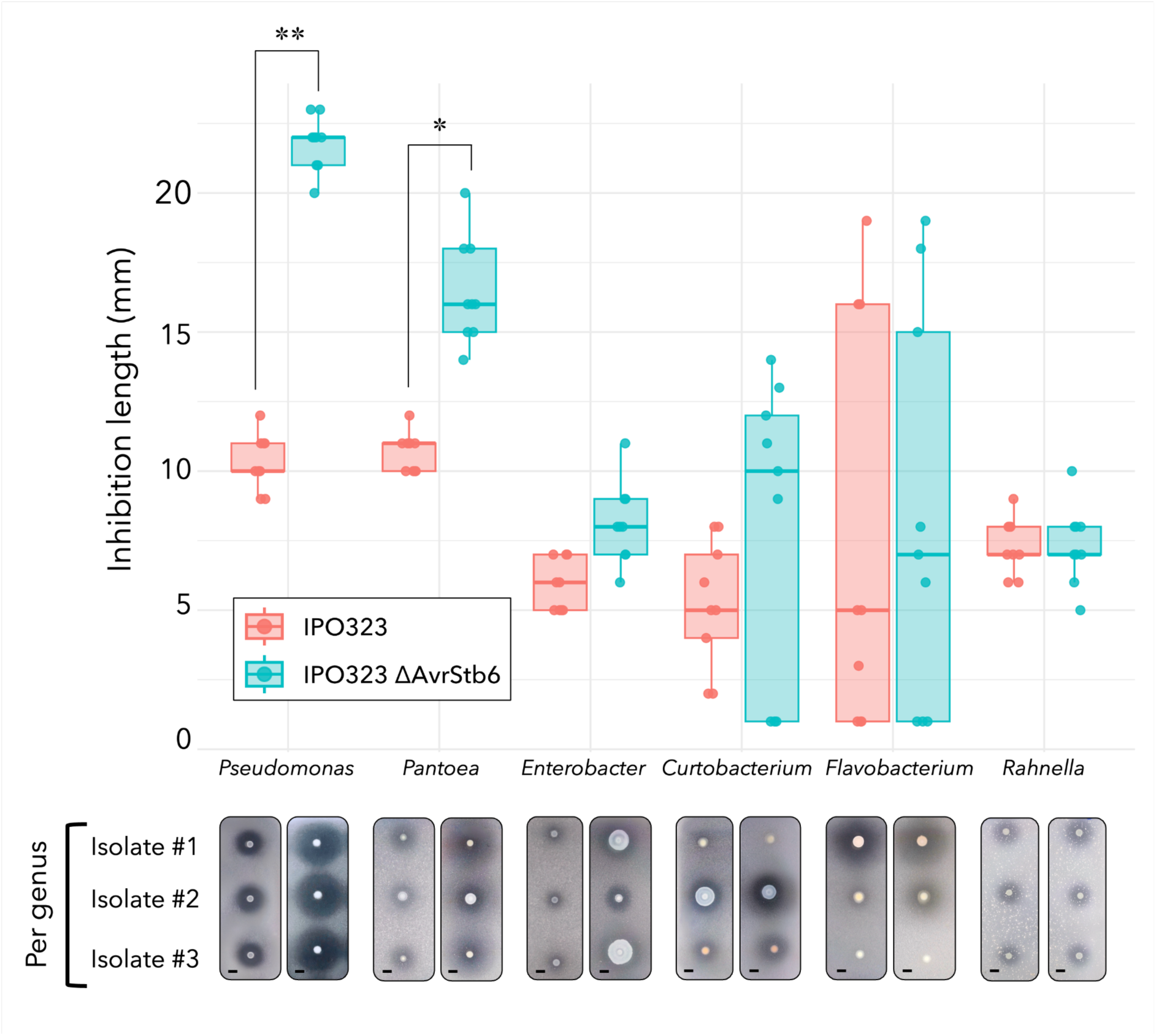
Differential inhibition of *Zymoseptoria tritici* strains by wheat-associated bacteria. Inhibition halo length produced by six bacterial genera against *Z. tritici* IPO323 (wild type) and IPO323 *ΔAvrStb6* mutant. Bacterial genera were selected based on differential relative abundance (S3 Fig), with three strains tested per genus (S2 Table). Box plots show median and interquartile range across; points represent individual measurements (n = 3 per isolate, n = 9 per genus). Asterisks indicate statistically significant differences between fungal strains based on unpaired t-tests (**p < 0.01, *p < 0.05). Representative confrontation assays are shown below, with fungal spores embedded in agar and bacterial inoculum applied on the surface. Scale bar: 10 mm.

Inhibitory effects varied among bacterial isolates, both within and between genera, and between fungal strains. Notably, *Pseudomonas* and *Pantoea* isolates exhibited strong and largely strain-independent inhibition of the IPO323 Δ*AvrStb6* mutant relative to the IPO323 wild type. In contrast, *Enterobacter*, *Curtobacterium*, and *Flavobacterium* isolates displayed more variable and strain-dependent effects, ranging from no detectable inhibition to stronger inhibition of the IPO323 Δ*AvrStb6* mutant (S4 Fig). *Rahnella* isolates showed comparable inhibition of both *Z. tritici* strains regardless of the isolate tested (Fig 4). IPO323 wild-type and Δ*AvrStb6* strains exhibited comparable growth under bacteria-free in vitro conditions (S2 Fig), indicating that the observed differences are not attributable to intrinsic fungal growth defects.

### AvrStb6-mediated protection against antagonistic bacteria is conserved across distinct allelic variants and genetic backgrounds

To assess whether bacterial protection is associated specifically with the avirulent Stb6-recognized *AvrStb6* allele (IPO323/1E4 allele) or is conserved across virulent allelic variants, we tested this phenotype in the Swiss *Z. tritici* isolate 3D7, which carries an *AvrStb6* allele that evades recognition by the wheat immune receptor Stb6 (Zhan et al., 2005; Stewart & McDonald, 2014; Alassimone et al., 2024). We used two genetically modified 3D7 strains: a strain expressing a cytosolic mCherry marker (3D7-mCherry) and an mCherry-labelled strain lacking *AvrStb6* (3D7-mCherry Δ*AvrStb6*). These strains were used in in vitro confrontation assays with wheat *Pseudomonas* and *Pantoea* isolates (Fig 5A).

**Figure 5.**
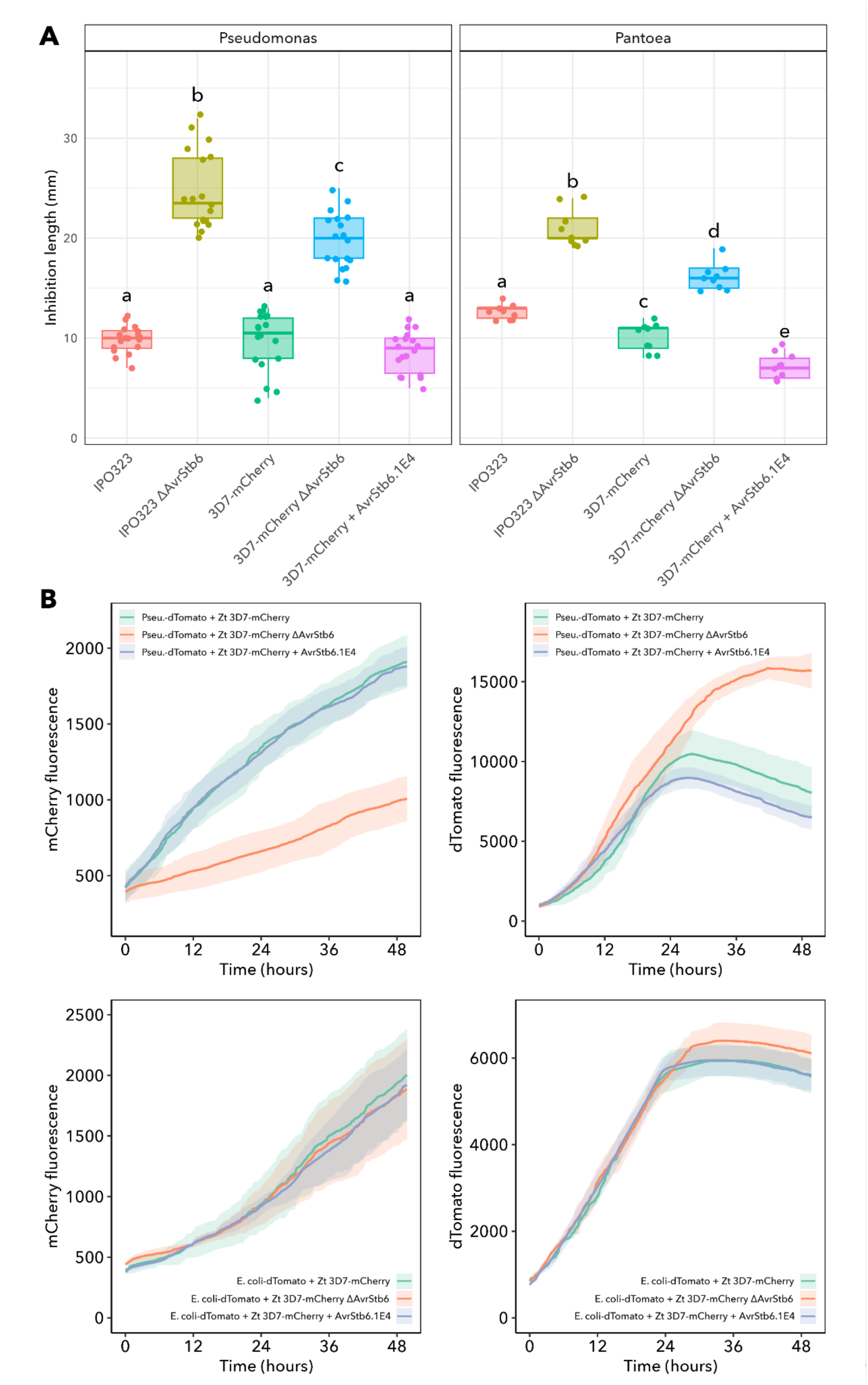
AvrStb6 confers protection against bacterial antagonism across genetic backgrounds and modulates fungal–bacterial dynamics. (A) Fungal inhibition caused by *Pseudomonas* and *Pantoea* co-cultivated with five *Z. tritici* strains: IPO323, IPO323 *ΔAvrStb6*, 3D7-mCherry, 3D7-mCherry *ΔAvrStb6*, and 3D7-mCherry AvrStb6_1E4_. Measures of inhibition represent halo lengths in fungal cultures. Box plots show median and interquartile range from independent biological replicates, with individual data points overlaid. Different letters indicate significant differences among *Z. tritici* isolates within each bacterial genus (one-way ANOVA followed by Tukey’s HSD, adjusted p < 0.05). (B) Time-course fluorescence measurements of co-cultures between dTomato-labelled *Pseudomonas* or *Escherichia coli* and mCherry-labelled *Z. tritici* 3D7 strains over 48 h. In co-culture with the *ΔAvrStb6* mutant, fungal mCherry fluorescence is reduced, while bacterial dTomato fluorescence is increased relative to co-cultures with AvrStb6-expressing strains (3D7-mCherry and 3D7-mCherry AvrStb6_1E4_). Shaded areas represent variation across replicates.

Consistent with the observations obtained for IPO323, deletion of *AvrStb6* in the 3D7 background resulted in stronger bacterial inhibition of fungal growth. Bacteria-mediated inhibition of 3D7-mCherry Δ*AvrStb6* was approximately two-fold higher during confrontation with *Pseudomonas* isolates and 1.5-fold higher during confrontation with *Pantoea* isolates compared with 3D7-mCherry (Fig 5A). In addition, we included a transgenic 3D7 strain expressing the avirulent *AvrStb6_1E4_* allele (3D7-mCherry + *AvrStb6_1E4_*) to assess whether introduction of a Stb6-recognized allele alters bacterial protection in the 3D7 background. Expression of *AvrStb6_1E4_* reduced sensitivity to *Pseudomonas* and *Pantoea* isolates relative to the 3D7-mCherry Δ*AvrStb6* strain and, in the case of *Pantoea* confrontations, resulted in even lower inhibition than observed for 3D7-mCherry (Fig 5A).

To further examine these interactions, mCherry-labelled 3D7 strains were co-cultured with dTomato-labelled wheat-associated *Pseudomonas* isolates and fluorescence was monitored over 48 h (Fig 5B). In the absence of *AvrStb6* (3D7-mCherry Δ*AvrStb6*), fungal mCherry fluorescence was reduced in the presence of *Pseudomonas*, whereas bacterial dTomato fluorescence increased relative to co-cultures containing *AvrStb6*-expressing strains (3D7-mCherry and 3D7-mCherry + *AvrStb6_1E4_*). This phenotype was not observed when mCherry-labelled 3D7 strains were co-cultured with dTomato-labelled *Escherichia coli* (Fig 5B), suggesting that the effects of *AvrStb6* are not broadly associated with bacterial interactions but may instead be specific to antagonistic wheat-associated bacteria such as *Pseudomonas* spp. Together, these results support a role for AvrStb6 in reducing the sensitivity of *Z. tritici* to selected wheat-associated antagonistic bacteria.

### AvrStb6 associates with the *Z. tritici* cell wall

Previous studies demonstrated that GFP-tagged AvrStb6_1E4_ is secreted extracellularly and potentially accumulates in developing apoplastic hyphae of *Z. tritici* (Alassimone et al., 2024). Given the predicted surface-binding properties of AvrStb6 (Fig 1B) and its role in reducing sensitivity to antagonistic bacteria (Fig 4 and 5), we hypothesized that AvrStb6 protects the fungal hyphae against antagonistic bacteria in the apoplast. We therefore set out to investigate whether AvrStb6 can associate with the fungal cell wall. To test this, 6xHis-tagged AvrStb6_1E4_ protein was incubated with cell wall extracts (alcohol-insoluble residue, AIR) prepared from 6-day-old hyphal cultures of 3D7-mCherry Δ*AvrStb6*. Following incubation, pellet and supernatant fractions were separated and analysed for the presence of 6xHis-AvrStb6_1E4_ by Western blot.

6xHis-AvrStb6_1E4_ was detected exclusively in the cell wall pellet fraction, with no detectable protein present in the supernatant (Fig 6), indicating that AvrStb6 associates with the *Z. tritici* cell wall in vitro. Similarly, the virulent variant 6xHis-AvrStb6_1A5_ was found predominantly in the cell wall pellet fraction, although a weak signal was observed in the supernatant (Fig 6). To determine whether this interaction reflects a general property of secreted fungal proteins, or an experimental outcome of the approach, we performed the same assay using two His-tagged secreted killer protein (KP)-like effectors from *Zymoseptoria passerinii* (KP4-like effector Zpa796_jg5892.t1 and KP-6 like effector Zpa796_jg2861.t1). In contrast to AvrStb6, neither KP-like effector showed predominant association with the *Z. tritici* cell wall (Fig 6, S6 Fig). 6xHis-Zpa796_jg5892.t1 was detected primarily in the supernatant fraction, with a weak signal detected in the pellet fraction, whereas 6xHis-Zpa796_jg2861.t1 was detected exclusively in the supernatant (Fig 6, S6 Fig). These results indicate that the specific association with the fungal cell wall is not a general feature of secreted *Zymoseptoria* effectors and support a specific interaction between virulent and avirulent AvrStb6 protein variants and the *Z. tritici* cell wall.

**Figure 6.**
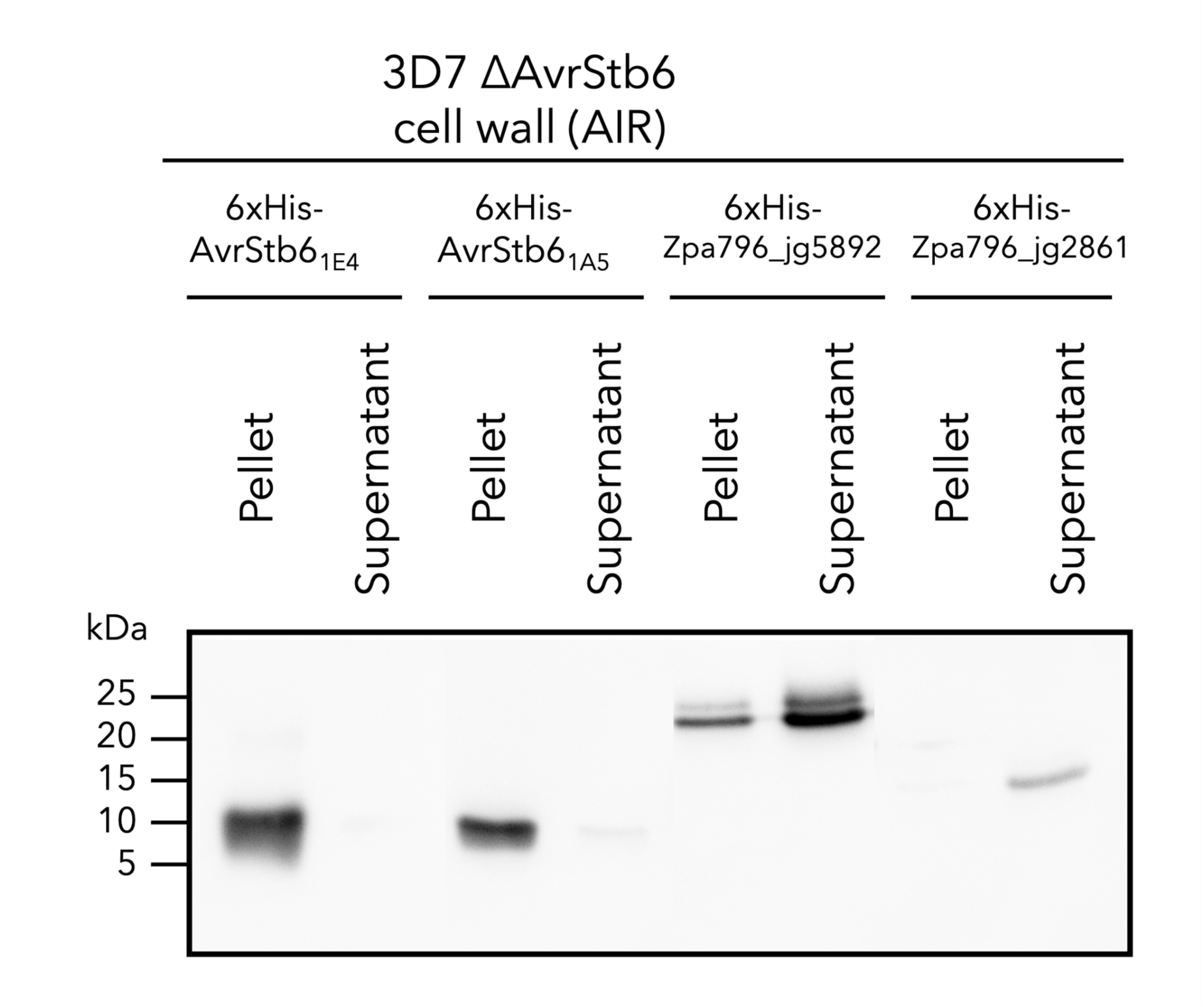
AvrStb6 isoforms associate with the *Zymoseptoria tritici* cell wall. Western blot analysis of 6xHis-tagged effector proteins following incubation with alcohol-insoluble residue (AIR) cell wall extracts obtained from 6-day-old hyphal cultures of the *Z. tritici* 3D7-mCherry Δ*AvrStb6* strain. After incubation for 1 h at 4°C, samples were separated into pellet (cell wall-associated fraction) and supernatant fractions prior to western blot analysis. The cell wall association comparison included two AvrStb6 protein variants (6xHis-AvrStb6_1E4_ and 6xHis-AvrStb6_1A5_) and two secreted killer protein (KP)-like effectors from *Zymoseptoria passerinii* (KP4-like effector Zpa796_jg5892.t1 and KP6-like effector Zpa796_jg2861.t1). AvrStb6 variants were predominantly detected in the pellet fraction, whereas KP-like effectors were detected mainly in the supernatant fraction. Corresponding control blots are shown in S6 Fig. Molecular mass marker positions (kDa) are indicated on the left.

## Conclusions

Collectively, our findings support a previously uncharacterized role for AvrStb6 in mediating interactions between *Z. tritici* and the wheat-associated microbiota. While fungal effectors are traditionally studied almost exclusively through the lens of plant immune recognition, our results suggest that AvrStb6 contributes to fungal fitness by reducing sensitivity to antagonistic bacteria during host colonization. Although computational analyses initially suggested that AvrStb6 may function as an antimicrobial protein, in vitro growth inhibition assays demonstrated no detectable antibacterial activity against a diverse panel of wheat-associated bacteria. Instead, deletion of *AvrStb6*, in two distinct genetic backgrounds of *Z. tritici*, consistently increased fungal sensitivity to antagonistic bacterial isolates, particularly *Pseudomonas* spp., across multiple fungal genetic backgrounds and across both virulent and avirulent AvrStb6 variants.

Importantly, we demonstrate that AvrStb6 associates with the *Z. tritici* cell wall in vitro, whereas unrelated secreted *Zymoseptoria* effectors do not display the same property. Together with previous studies showing extracellular secretion and accumulation of AvrStb6 in developing fungal hyphae, these findings support a model in which AvrStb6 reassociates with the fungal surface to protect *Z. tritici* against antagonistic members of the wheat microbiota. This work expands the current view of fungal effectors beyond their established roles in host immune manipulation and suggests that selective pressures imposed by microbial competition may also contribute to the long-term maintenance and diversification of avirulence effector genes in pathogen populations.

## Data Availability

Raw amplicon sequence data: BioProject: PRJNA1474436 RStudio data analysis: https://github.com/liz-florez/AvrStb6-wheat_microbiome_interactions

## Authors contribution

LF: Conceptualization, Methodology, Data Curation, Formal Analysis, Investigation, Supervision, Validation, Visualization, Writing - Original Draft Preparation

VF: Data Curation, Formal Analysis, Writing – Review & Editing. HB: Investigation, Methodology, Writing – Review & Editing.

ML: Investigation, Writing – Review & Editing. LC: Data Curation, Software

AT: Data Curation, Software

CF: Conceptualization, Investigation, Methodology, Resources SB: Methodology, Resources

GR: Methodology, Resources

ASV: Resources, Writing – Review & Editing.

ES: Conceptualization, Funding Acquisition, Supervision, Writing – Original Draft Preparation, Writing – Review & Editing.

## Acknowledgements

This work was funded by the German Science Foundation (DFG) in the CRC1182 program, and through the ERC Consolidator grant 101087809 FungalSecrets awarded to EHS. The authors wish to thank members of the Environmental Genomics group for helpful discussions, Elisha Thynne for advice and suggestions and Nadja Hammelmann for support with the initial experimental set up.

## Supplementary Figures

**Supplementary Figure 1.**
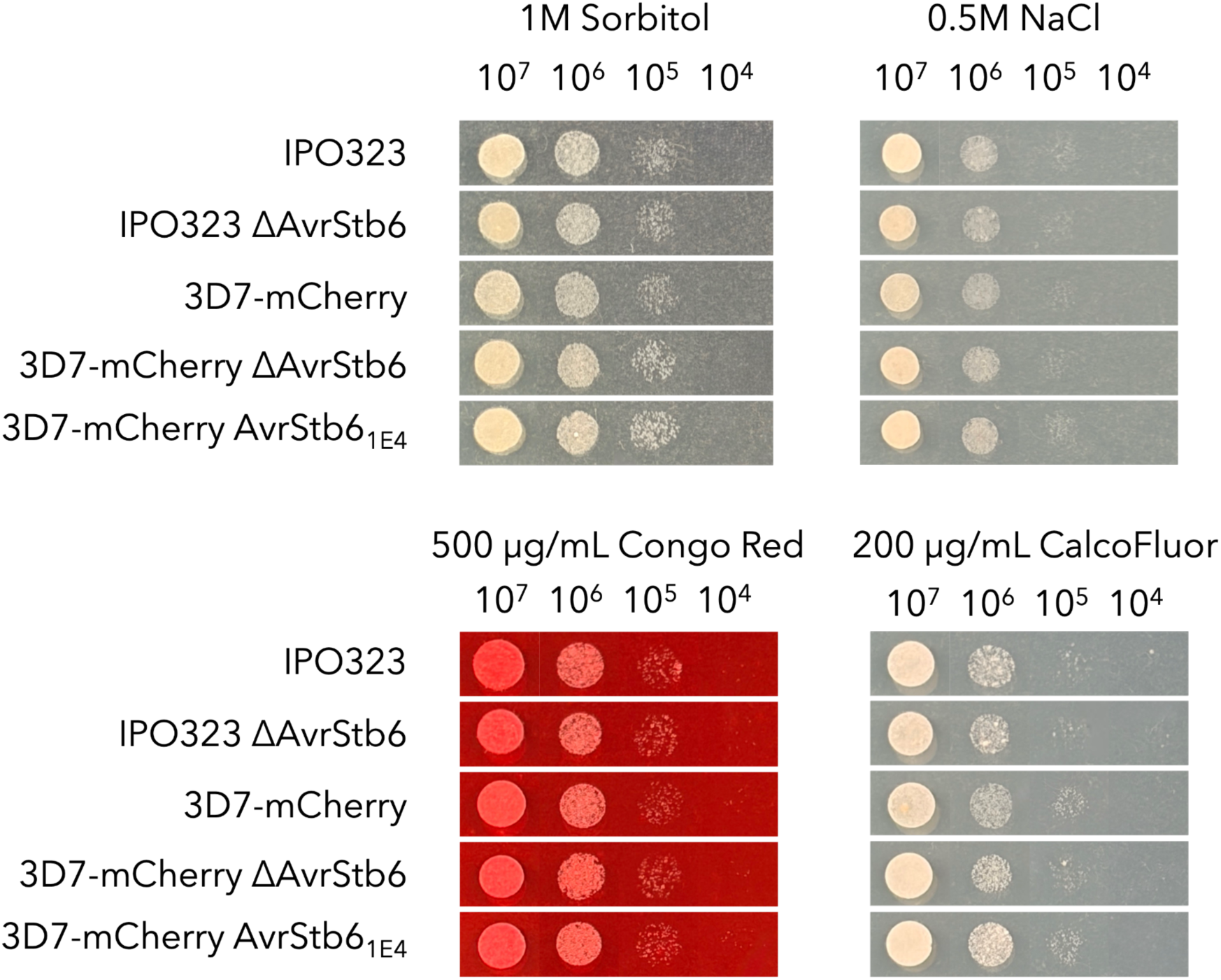
*In vitro* stress assay and colony morphology of *Zymoseptoria tritici* wild-type and mutant lines. Growth of *Z. tritici* IPO323 and 3D7 lines was tested under osmotic stress (1M Sorbitol, 0.5M NaCl) and cell wall stress (500 µg/mL Congo Red, 200µg/mL CalcoFluor). The phenotypes were assessed at four days post incubation to mimic the same timeline followed for the *in vitro* confrontation assay. No growth-or colony morphology-phenotypic differences were observed across the strains.

**Supplementary Figure 2.**
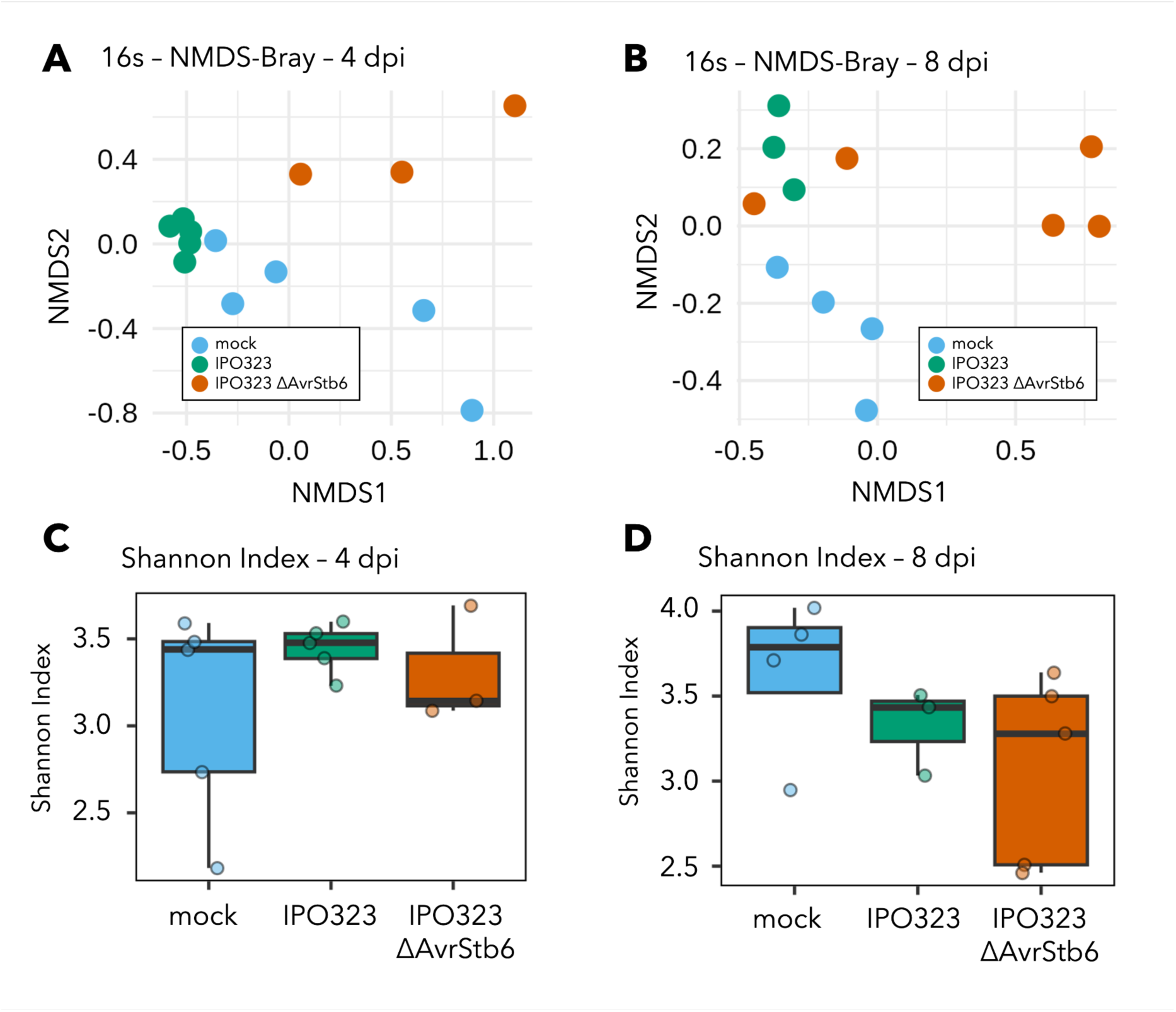
Bacterial community structure and alpha diversity across treatments. (A–B) Non-metric multidimensional scaling (NMDS) ordination based on Bray–Curtis dissimilarities of 16S-derived bacterial communities at 4 and 8 days post inoculation (dpi). Each point represents a sample, and distances between points reflect differences in community composition. At 4 dpi, samples cluster by treatment, with IPO323-associated communities separating from mock, while ΔAvrStb6 shows a distinct distribution. At 8 dpi, separation between treatments remains but appears reduced. (C–D) Shannon diversity index of bacterial communities at 4 and 8 dpi. Boxes indicate median and interquartile range, with points representing individual samples. Diversity is comparable across treatments at 4 dpi, whereas at 8 dpi mock samples show higher diversity relative to infected treatments, with ΔAvrStb6 tending toward lower values.

**Supplementary Fig. 3.**
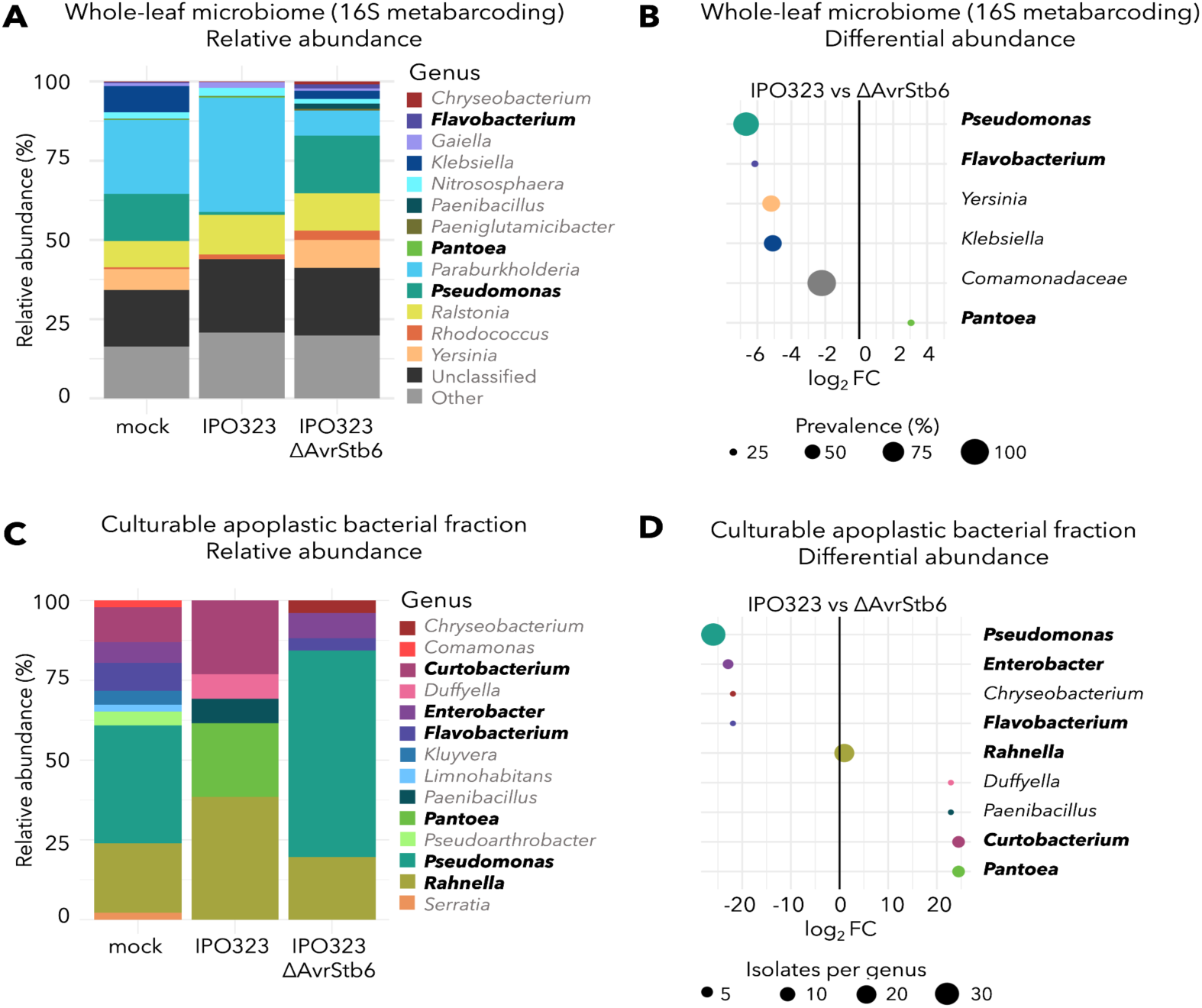
Comparison between whole-leaf microbiome profiling and culturable apoplastic bacterial isolates from wheat associated with *Zymoseptoria tritici* infection in wheat. (A) Relative abundance of bacterial genera detected by 16S rRNA metabarcoding of whole wheat leaves infected with mock, IPO323, or IPO323 Δ*AvrStb6* at 4 dpi. Relative abundance values are shown at the genus level. (B) Differential abundance analysis of bacterial genera identified in whole-leaf microbiome samples comparing IPO323 and IPO323 Δ*AvrStb6* treatments. Bubble size indicates prevalence across samples. Positive log2FC values indicate higher relative abundance in IPO323, whereas negative values indicate higher relative abundance in IPO323 Δ*AvrStb6*. (C) Relative abundance of culturable bacterial genera isolated from wheat apoplastic fluid collected from mock-, IPO323-, and IPO323 Δ*AvrStb6*-infected leaves. Relative abundance values were calculated based on the number of isolates assigned to each genus following partial 16S rRNA gene amplicon sequencing. (D) Differential abundance analysis of bacterial genera recovered from the culturable apoplastic fraction comparing IPO323 and IPO323 Δ*AvrStb6* treatments. Bubble size indicates the number of isolates recovered per genus. Positive log2FC values indicate higher relative abundance in IPO323, whereas negative values indicate higher relative abundance in IPO323 Δ*AvrStb6*. Genera selected for antimicrobial and confrontation assays are highlighted in bold.

**Supplementary Figure 4.**
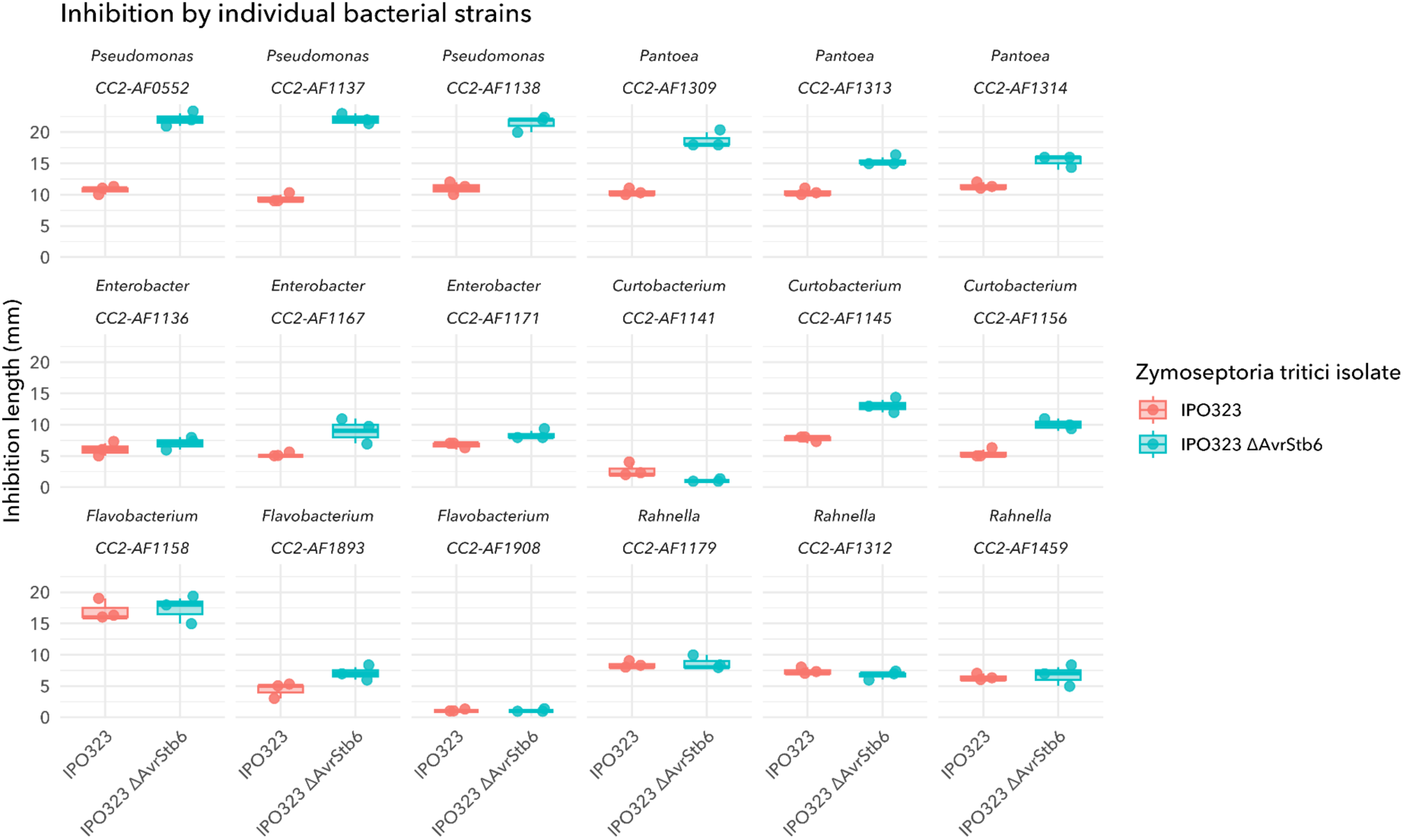
Strain-level variation in bacterial inhibition of *Zymoseptoria tritici*. Inhibition halo length produced by individual bacterial strains against *Z. tritici* IPO323 (wild type) and ΔAvrStb6. Each panel represents a single bacterial strain (CC2-AF ID) grouped by genus. Points show individual biological replicates (n = 3), and box plots indicate median and interquartile range. This representation highlights consistent responses across strains for some genera (e.g. *Pseudomonas*, *Pantoea*), whereas others display substantial intra-genus variability (e.g. *Curtobacterium*).

**Supplementary Figure 5.**
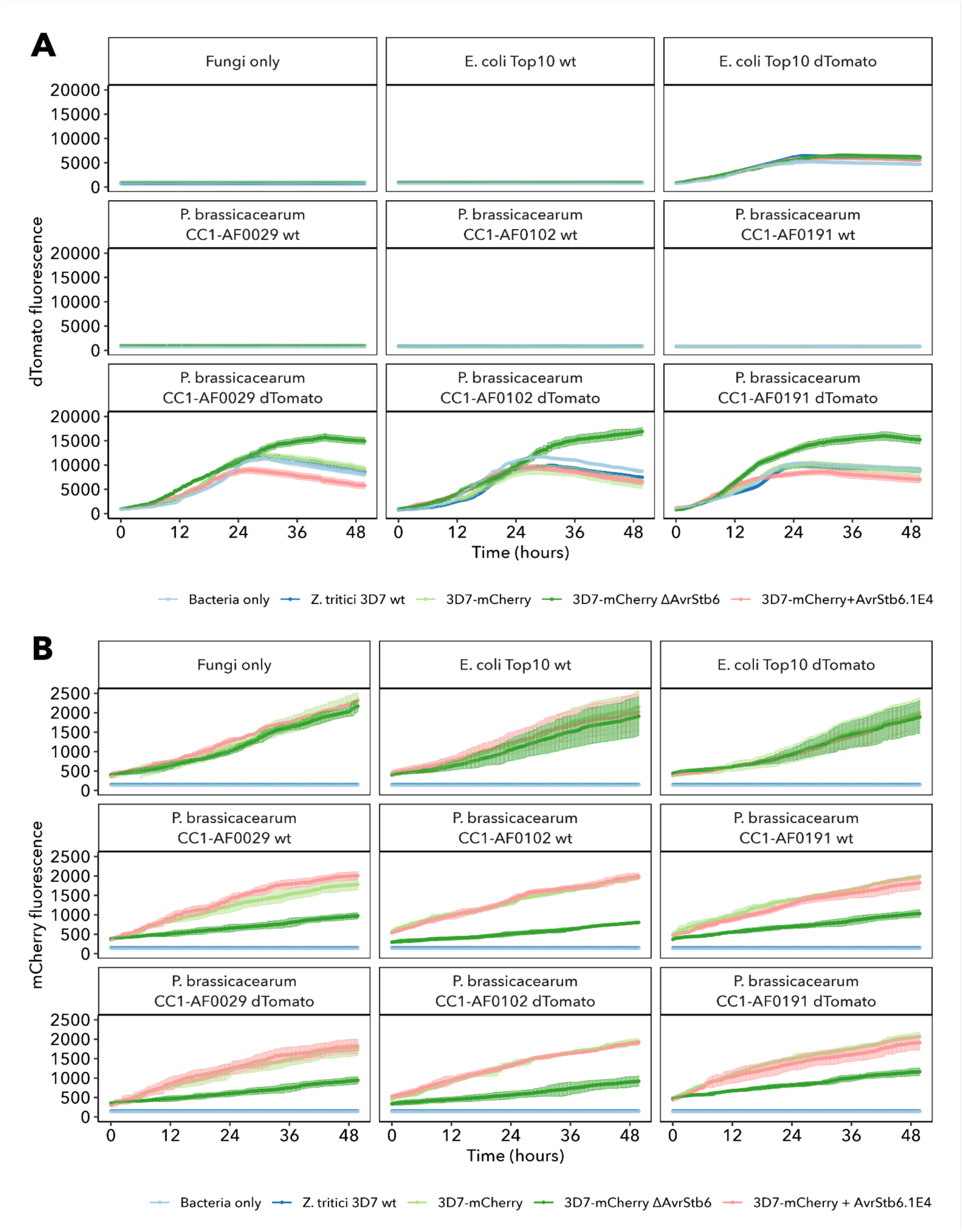
Validation controls and strain-specific fluorescence dynamics in *Zymoseptoria tritici*–*Pseudomonas brassicacearum* co-culture assays. (A) dTomato fluorescence measurements of co-cultures between individual *P. brassicacearum* isolates and *Z. tritici* strains over 48 h. Wells containing only fungal strains, non-fluorescent bacterial strains, or dTomato-labelled bacterial strains were included as fluorescence controls. (B) mCherry fluorescence measurements of the corresponding co-cultures shown in panel A. Shaded areas represent variation across replicates.

**Supplementary Figure 6.**
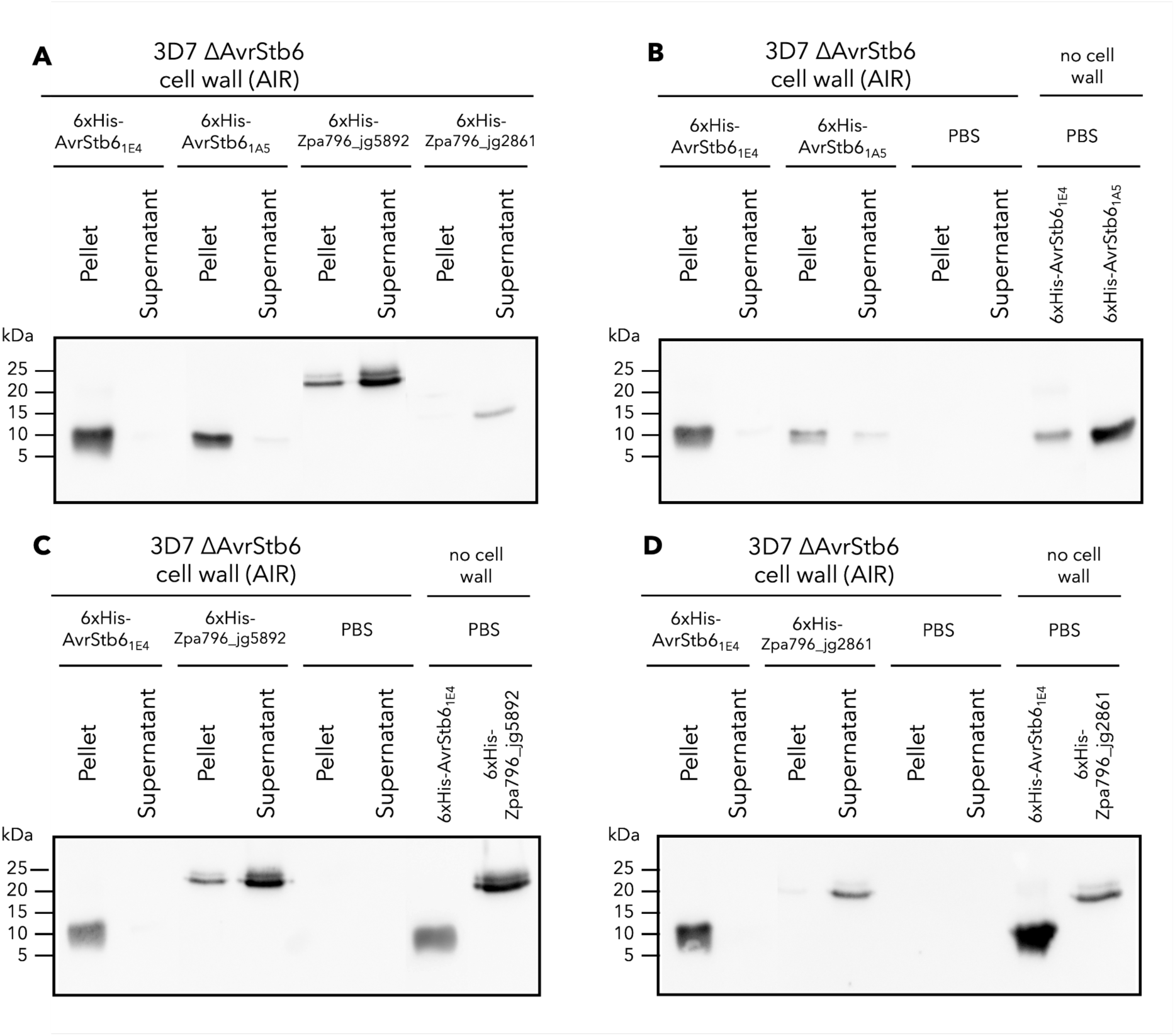
Control blots for the cell wall association assay shown in Figure 6. Cell wall association assay reproduced from Figure 6 for reference. (B–D) Control blots for individual proteins. AIR-only controls were incubated with PBS in the absence of recombinant protein, whereas protein-only controls were incubated in the absence of fungal cell wall extract. These controls confirmed the absence of detectable non-specific signals and verified the expected migration of each recombinant protein. Molecular mass marker positions (kDa) are indicated on the left.

## Supplementary Tables

**Supplementary Table 1.** *Zymoseptoria tritici* isolates used in this study.

| Fungal isolate | Origin | Description | Reference/Source |
| --- | --- | --- | --- |
| IPO323 | Netherlands | *Z. tritici* wild-type isolate carrying the avirulent *AvrStb6* allele recognized by wheat *Stb6*. | Stukenbrock lab |
| IPO323 Δ*AvrStb6* | Netherlands, genetically modified | IPO323 deletion mutant lacking *AvrStb6*. | Kema et al, 2018 |
| 3D7-mCherry | Switzerland, genetically modified | *Z. tritici* 3D7 strain carrying a virulent *AvrStb6* allele that evades recognition by wheat *Stb6* and constitutively expressing cytosolic mCherry. | Alassimone et al, 2024 |
| 3D7-mCherry Δ*AvrStb6* | Switzerland, genetically modified | 3D7-mCherry deletion mutant lacking *AvrStb6*<sub>3D7</sub> | Alassimone et al, 2024 |
| 3D7-mCherry + *AvrStb6*<sub>1E4</sub> | Switzerland, genetically modified | 3D7-mCherry strain ectopically expressing the avirulent *AvrStb6*<sub>1E4</sub> allele. | Alassimone et al, 2024 |

**Supplementary Table 2.**
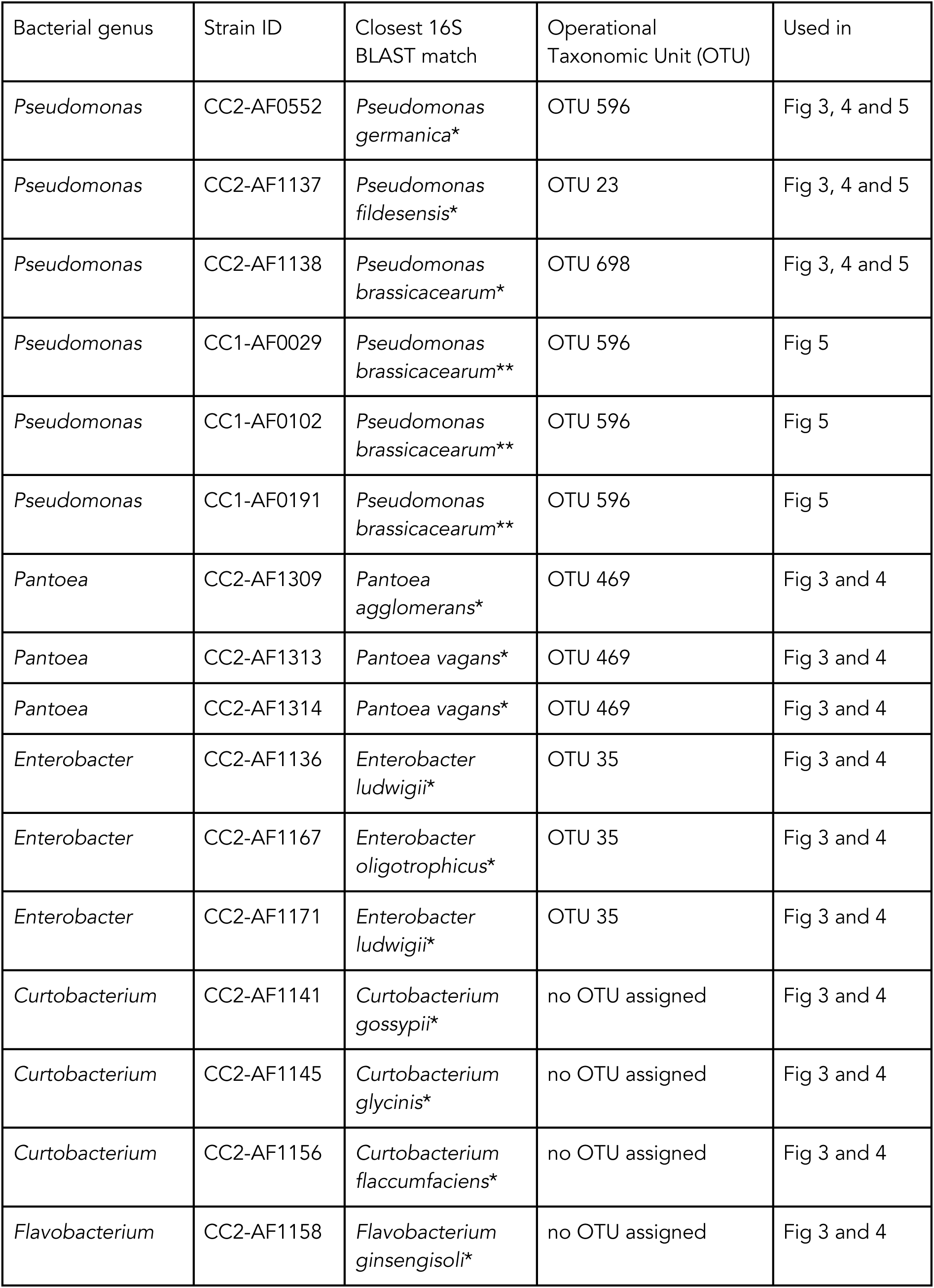
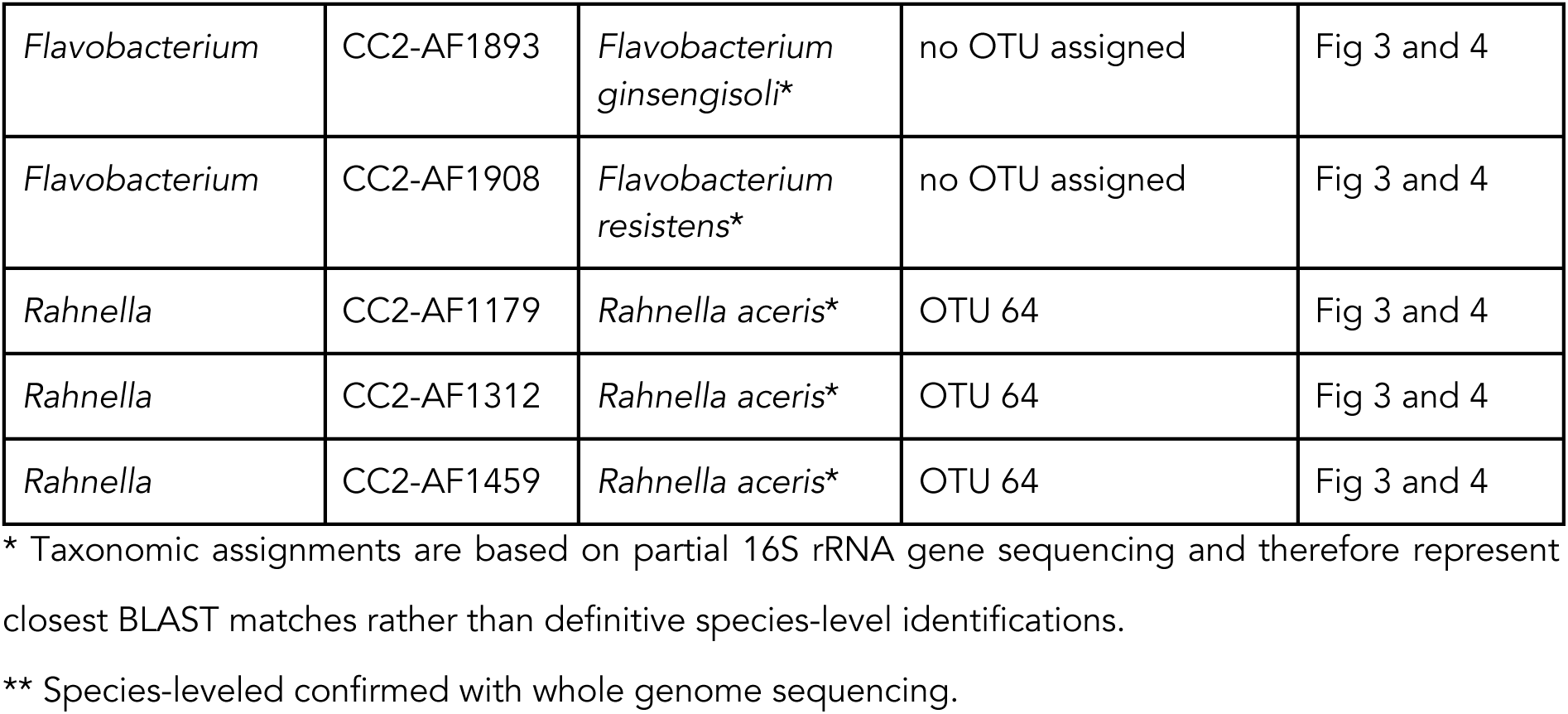
Wheat-associated bacterial isolates used in this study. All species were isolated from apoplastic fluids of *Triticum aestivum* cultivar Obelisk.

**Supplementary Table 3.**
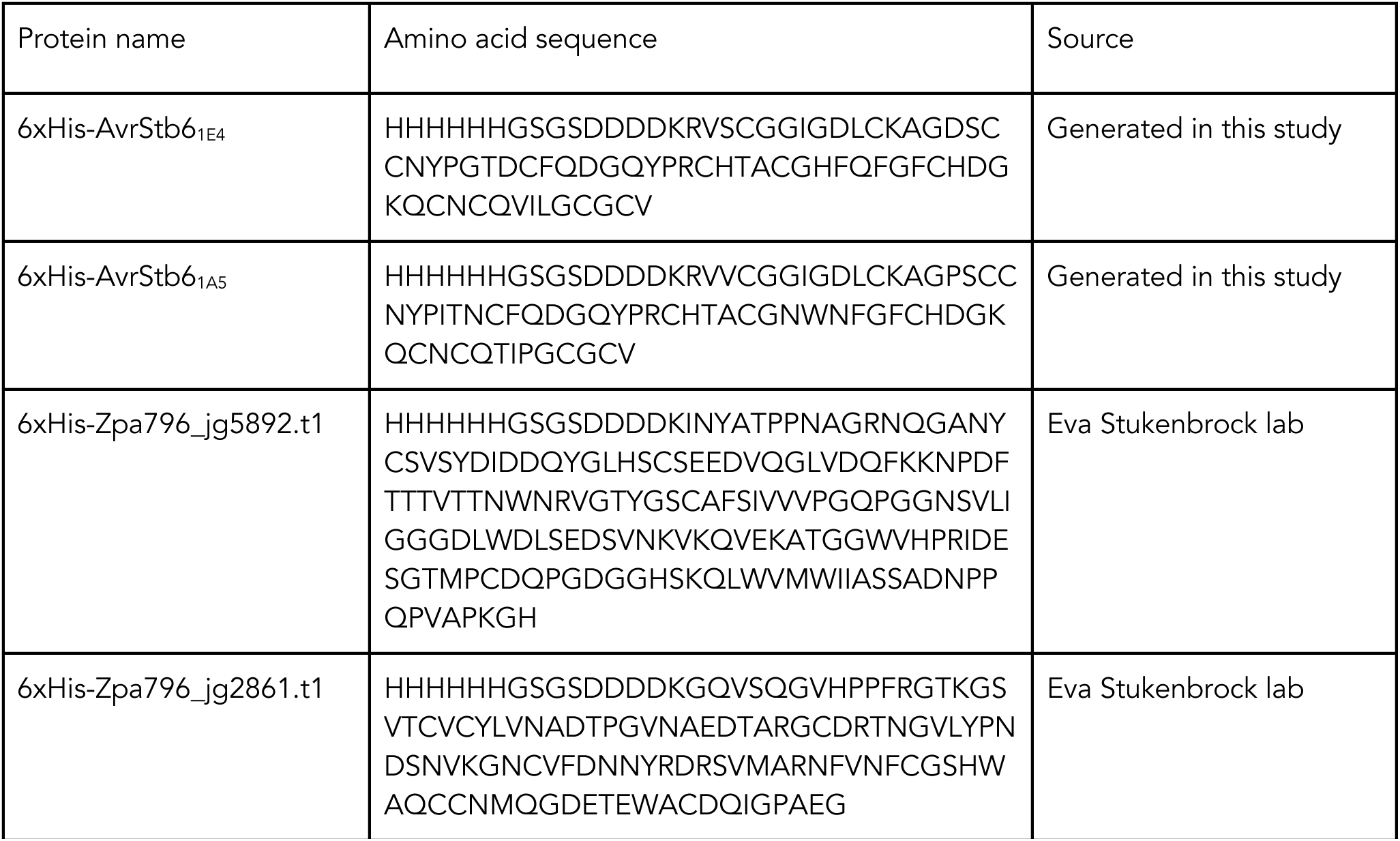
6xHis-tagged proteins used in this study. Amino acid sequence and source details are provided.

