## Supplementary figures and images for "An effector protein that protects a fungal pathogen from the plant microbiota during host colonisation"

### bacteria_vs_zymo_per_strain_S4_Fig.png

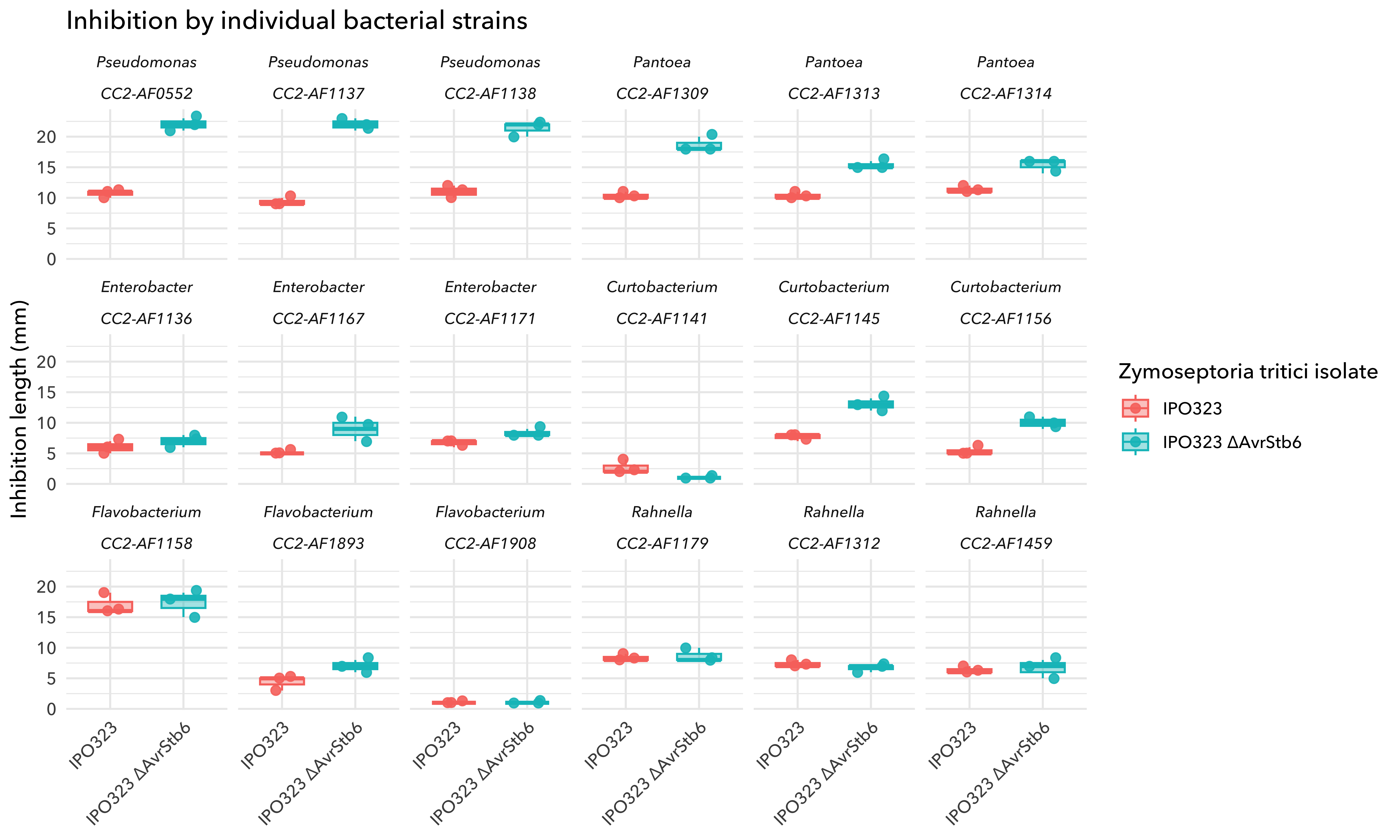

### Cell-wall_binding_S6_Fig.png

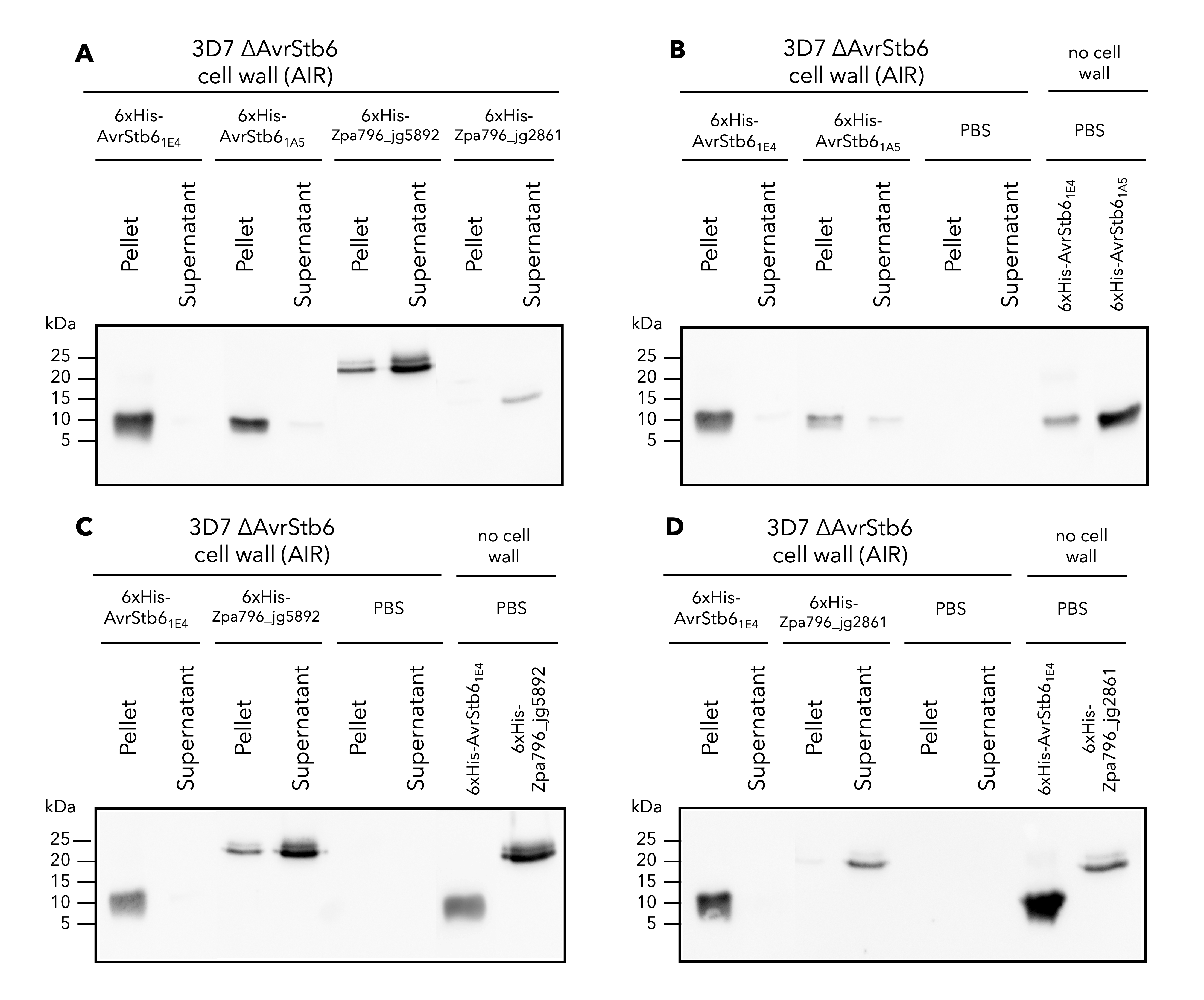

### Microbiome_shannon_NMDS_S2_Fig.png

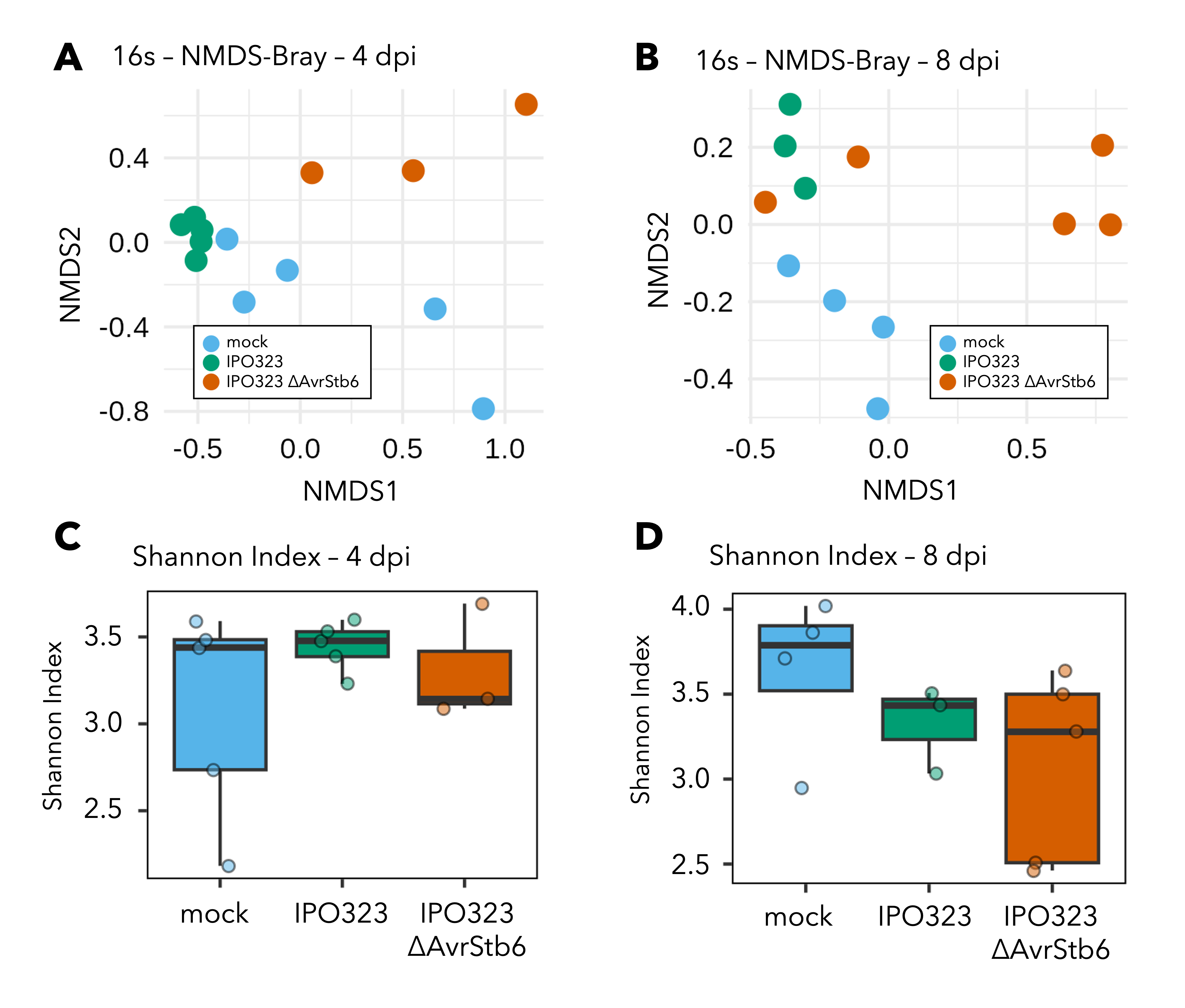

### RA_and_DA_plots_S3_Fig.png

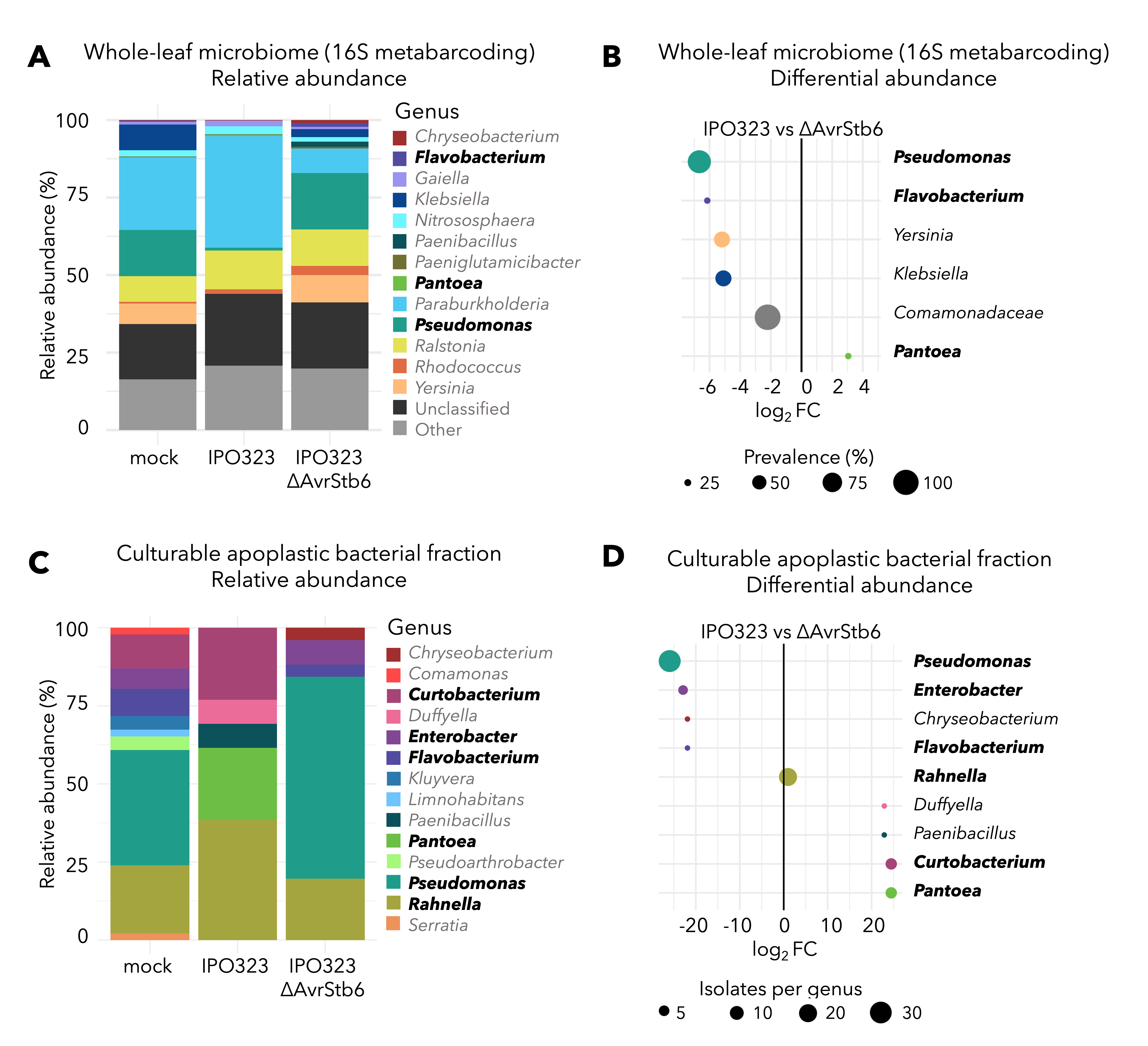

### Zymo_3D7_fluorescence_S5_Fig.png

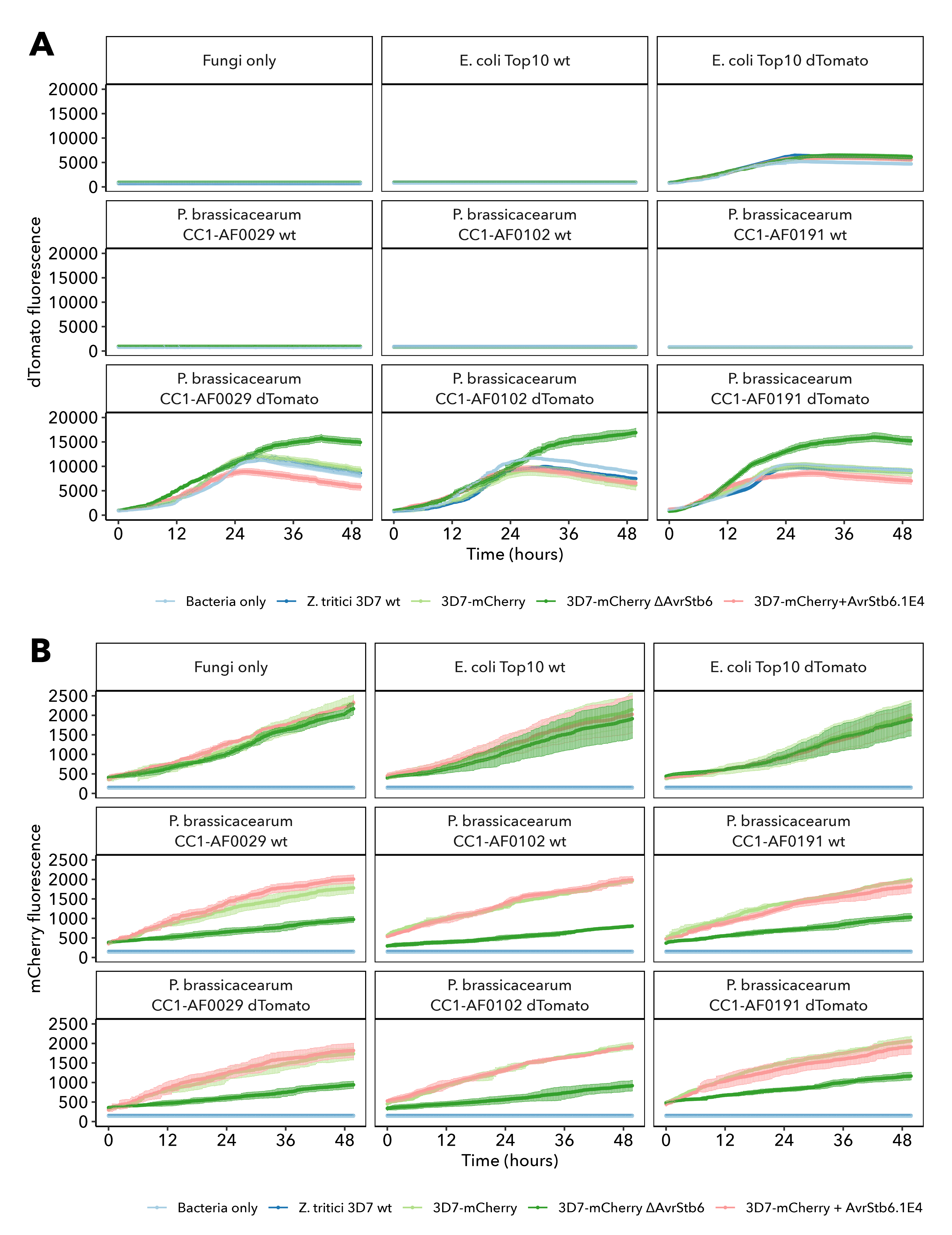

### Zymo_mutants_stress_assay_S1_Fig.png

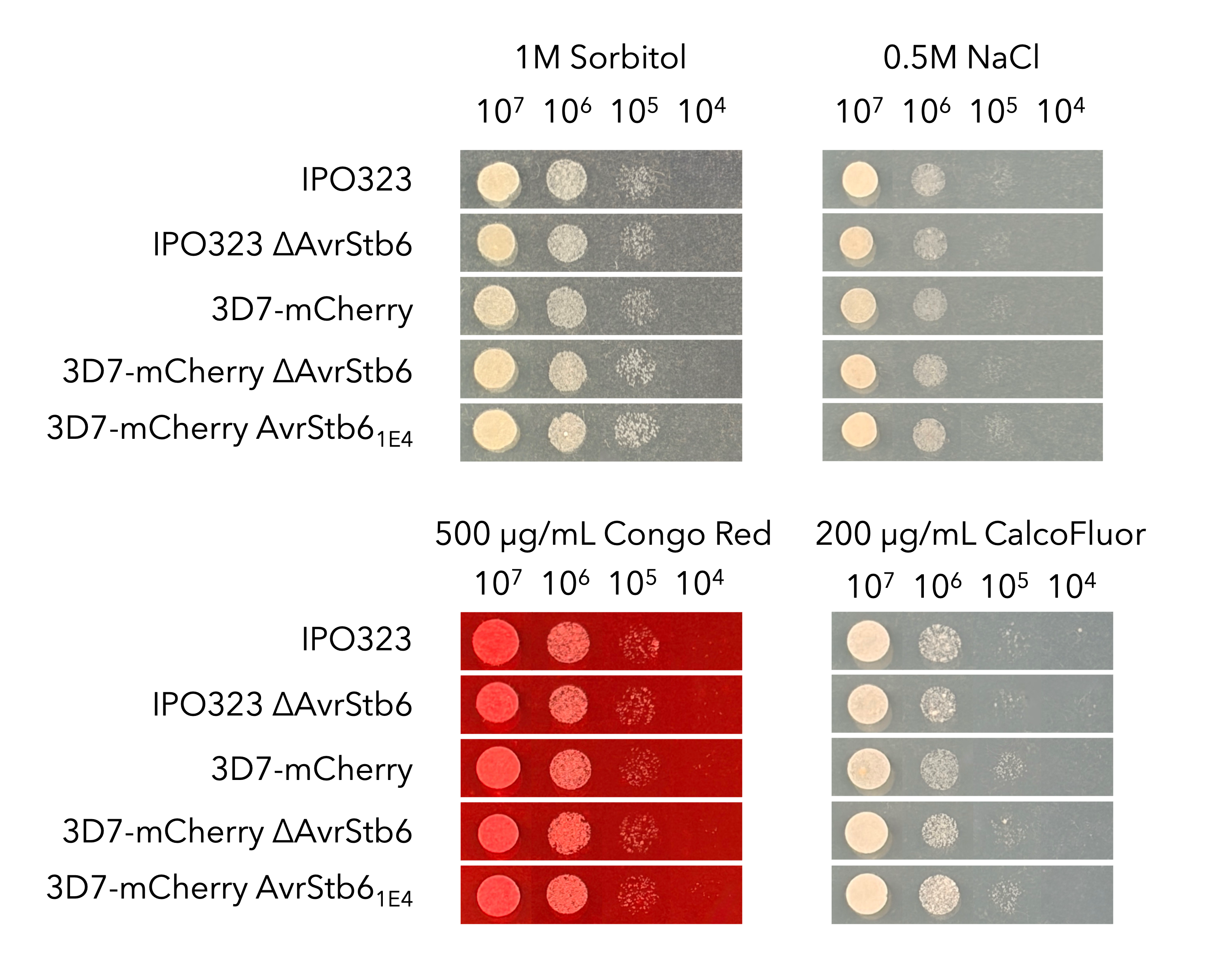
